# Compression-enabled interpretability of voxelwise encoding models

**DOI:** 10.1101/2022.07.09.494042

**Authors:** Fatemeh Kamali, Amir Abolfazl Suratgar, Mohammadbagher Menhaj, Reza Abbasi-Asl

## Abstract

Voxelwise encoding models based on convolutional neural networks (CNNs) have emerged as state-of-the-art predictive models of brain activity evoked by natural movies. Despite their superior predictive performance, the huge number of parameters in CNN-based models have made them difficult to interpret. Here, we investigate whether model compression can build more interpretable and more stable CNN-based voxelwise models while maintaining accuracy. We used multiple compression techniques to prune less important CNN filters and connections, a receptive field compression method to select receptive fields with optimal center and size, and principal component analysis to reduce dimensionality. We demonstrate that the model compression improves the accuracy of identifying visual stimuli in a hold-out test set. Additionally, compressed models offer a more stable interpretation of voxelwise pattern selectivity than uncompressed models. Finally, the receptive field-compressed models reveal that the optimal model-based population receptive fields become larger and more centralized along the ventral visual pathway. Overall, our findings support using model compression to build more interpretable voxelwise models.

## 1. Introduction

A prominent question in computational neuroscience is how sensory information is represented and processed in the visual cortex [1, 2] and various computational models have been developed to predict brain activity during sensory tasks. Constructing accurate and data-driven models requires large-scale data collection of the brain’s high-dimensional and complex activity. In the past decade, functional magnetic resonance imaging (fMRI) has emerged as a standard technique to record brain activity during natural visual tasks [1–7], and computational models of fMRI blood oxygen level-dependent (BOLD) signals have been developed. Voxelwise encoding models of BOLD signals use sensory stimuli to predict brain activity during various visual tasks and are the focus of this study. On the other hand, decoding models use brain activity to reconstruct and categorize visual stimuli [2, 6, 8]. Together, these encoding and decoding models provide functional descriptions of cortical areas.

Voxelwise encoding models of BOLD signals are often composed of two components: (1) a feature extraction module to construct a rich feature set from visual stimulus, and (2) a response prediction module to accurately predict BOLD signals from stimulus features (**Figure 1A**). The feature extraction module has historically been built based on classical machine learning techniques such as local binary pattern (LBP) [9], fisher vector [3], word to vector [10], or Gabor wavelets [2, 11]. Recently, however, deep networks and, specifically, convolutional neural networks (CNNs), have emerged as the state-of-the-art encoding models of BOLD signals [8, 12–18]. These networks extract both high-level and low-level features through their hierarchical structure. For example, Agrawal et al. proposed a deep CNN to predict BOLD signals in the visual cortex and showed superior performance of CNN-based models compared to scale-invariant feature transform or SIFT-based models [3]. Other types of deep neural networks, including recurrent neural networks [19], autoencoders [5], deep residual networks [20], image captioning models [4], and capsule networks [21], have also offered accurate predictive models of human visual cortex responses. For the response prediction module, regularized linear regression has been the standard approach [3,7,8,15,19,20]. A linear model provides a simple map between non-linear features and the BOLD signal and therefore is more interpretable [7].

**Figure 1:**
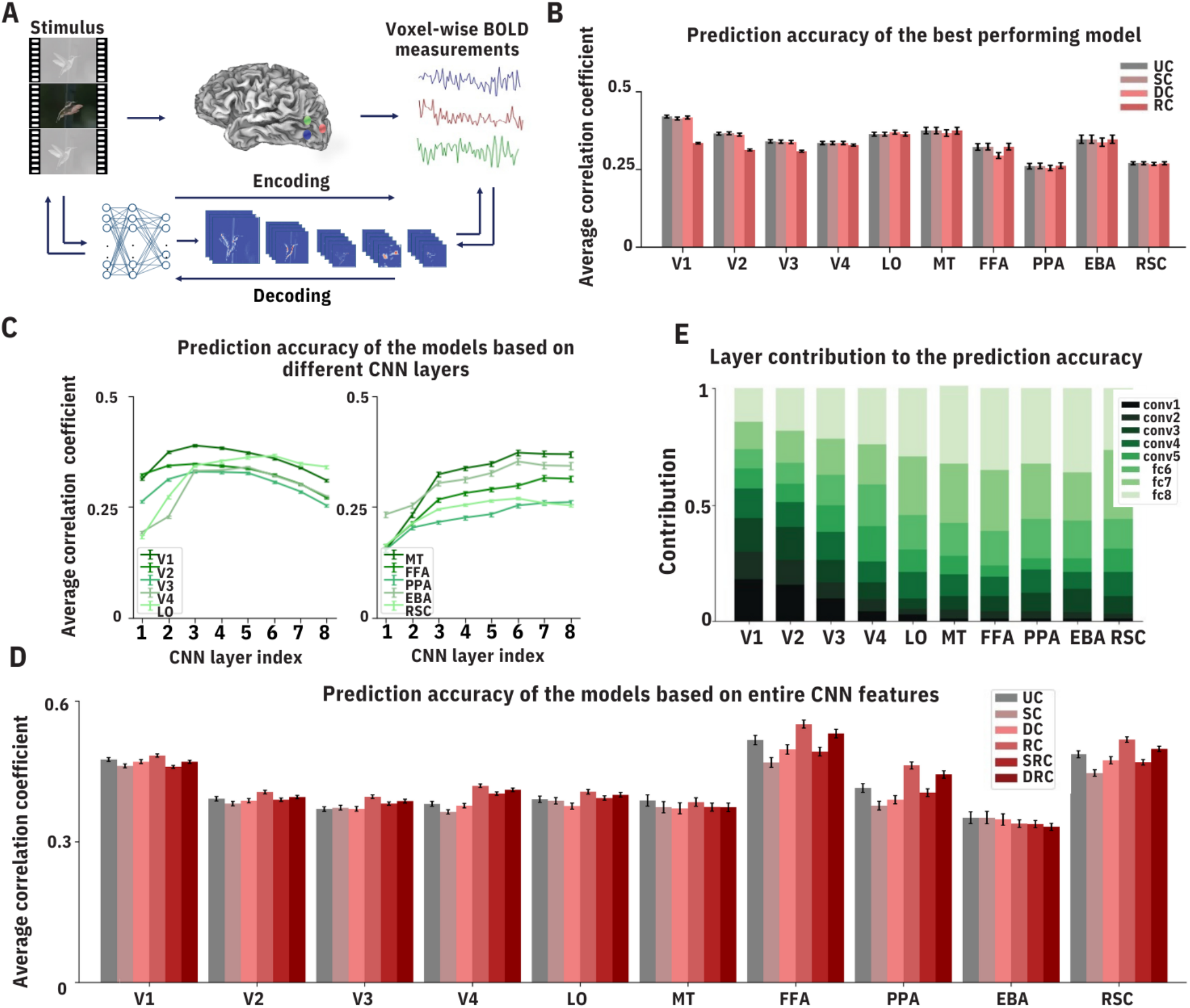
Compressed voxelwise encoding models accurately predict BOLD responses. **A**. An encoding model predicts the fMRI BOLD responses from the visual stimulus features. The decoding model predicts the optimal stimulus from the BOLD responses. **B**. The Pearson correlation coefficient between estimated response using the best CNN layer and measured fMRI response. The values are averaged over all voxels in each visual area across all subjects in both PURR and Vim-2 datasets and visualized for uncompressed (UC), structurally-compressed (SC), deep-compressed (DC), and receptive field-compressed (RC) models. Error bars show the 95% confidence interval. Compression does not significantly affect the accuracy. **C**. The Pearson correlation between estimated and measured fMRI responses for different CNN layers. Each data point shows the averaged correlation coefficient over all voxels in each visual area for the compressed model. Error bars show the 95% confidence interval. The contribution of the higher CNN layers is attenuated for early visual areas, while the reverse trend is visible for higher visual areas. **D**. Each bar indicates the Pearson correlation coefficient between estimated and measured fMRI response averaged over all voxels in a visual area and across all subjects in both PURR and Vim-2 datasets. The average correlation coefficients are shown for uncompressed (UC), structurally-compressed (SC), deep-compressed (DC) and receptive field-compressed (RC), structurally receptive field-compressed (SRC), and deep receptive field-compressed (DRC) models. Error bars show the 95% confidence interval. Compression does not significantly affect the accuracy. **E**. Each column indicates the contribution of each CNN layer to prediction accuracy. Feature maps extracted from lower CNN layers have a higher contribution to lower visual areas and feature maps extracted from higher CNN layers have a higher contribution to higher visual areas.

Despite their promising performance, models based on deep CNNs are often extremely hard to interpret [22–24]. Specifically, millions of parameters and the highly non-linear transformations in these models make them impossible for human observers and domain experts to understand. In many scientific applications, such as computational neuroscience, this form of post-hoc interpretation is essential in understanding the scientific phenomena underlying the model. Recently, model compression has emerged as an efficient technique for interpreting CNN-based models [25, 26]. Compression removes redundant components of the model while preserving the accuracy of the model; therefore, a compressed model is easier for a domain expert to understand. Additionally, a compressed model requires considerably fewer computational operations, and thus, is faster than the uncompressed model. This reduced computational cost could facilitate the application of compressed models in real-time prediction.

In this paper, we explore the role of model compression in building more interpretable CNN-based models of BOLD responses. We use multiple compression techniques applied to both the feature extraction module and the response prediction module. These compression techniques include (1) a recently established structural compression [25] method to prune less important CNN filters, (2) deep Compression [27] to remove less important connections, (3) a receptive field compression [16] to choose the receptive fields with the optimal center and size, and (4) principal component analysis (PCA) [28] to reduce the dimensionality of the model. Using two separate fMRI datasets collected during natural vision from 6 participants, we first demonstrate that the compression of CNN-based voxelwise models reduces their size and computational cost while maintaining their high predictive accuracy. We then show compression improves the accuracy of identifying visual stimuli from the BOLD signal in the hold-out test set. We establish that the compressed encoding models reveal increased category-selectivity along the ventral visual pathway with higher stability compared to uncompressed models. Finally, we leverage the compressed models to quantitatively compare the model-based population receptive field sizes and locations along the different visual pathways.

## 2. Methods

### 2.1. Datasets

To build and investigate the compressed voxelwise models, we used two separate fMRI BOLD signal datasets. Each dataset contains BOLD fMRI recordings from three healthy participants watching hours of natural movie clips. For each subject, the dataset is divided into non-overlapping training and test sets. Further information about each dataset is provided below.

#### 2.1.1. PURR Dataset

fMRI data were obtained from three healthy volunteers in a 3T MRI system with a temporal resolution of 2s. The training set contains 374 movie clips (continuous with a frame rate of 30 fps) in a 2.4-h movie, divided randomly into 18 8-minute sections; the test set contains 598 movie clips in a 40-min movie, divided randomly into 5 sections of 8-minutes and 24s each. The training set was repeated two times whereas the test set was repeated ten times. During each section, an 8-minute single video segment was shown; the first and the last movie frames were shown as a static picture for 12s. Stimuli were chosen from video blocks and YouTube. The fMRI data were preprocessed and co-registered onto a standard cortical surface template using the processing pipeline for the Human Connectome Project [29]. The visual areas were defined with multi-modal cortical parcellation [30]. Additional details on this dataset can be found in [8].

#### 2.1.2. Vim-2 Dataset

fMRI data were obtained from three healthy volunteers in a 4T MRI system with a temporal resolution of 1s [31]. The training set includes a 2-hour movie; the test set includes a 9-minute movie. The training set was shown only once whereas the test set was repeated ten times. Stimuli were chosen from Apple Quick-Time HD gallery and YouTube. Retinotopic mapping data collected from the same subjects in separate scan sessions was used to assign voxels to visual areas [32]. This dataset is further described in [33].

### 2.2. Feature extraction *via* Convolutional Neural Networks

Our voxelwise encoding models consist of two modules: 1) The CNN-based feature extraction module which constructs a feature set for each frame in the visual stimulus. 2) the response prediction module which predicts the BOLD signal from the CNN-based features.

To extract features from each frame in the visual stimulus, we used AlexNet [34], a well-known CNN model pre-trained on the ImageNet dataset [34]. AlexNet consists of eight layers– the first five layers are convolutional and the last three are fully connected. The five convolutional layers use rectified linear activation functions, with the first layer receiving a 227 × 227 input image. Max-pooling is applied between layer 1 and layer 2, between layer 2 and layer 3, and between layer 5 and layer 6. The last layer uses a softmax function to generate a probability vector, from which an input stimulus frame is classified into 1000 classes. Layer 1 through layer 5 contain 96 kernels of 11 × 11, 256 kernels of 27 × 227, 384 kernels of 13 × 13, 384 kernels of 13 × 13, and 256 kernels of 13 × 13, respectively. Layers 6 to 8 contain 4096, 4096, and 1000 units, respectively.

For the response prediction module, we used a linear regression model with *L*_2_-norm regularization (Ridge regression [35]). We used each CNN layer output to predict the voxelwise responses and then selected the layer with the highest accuracy for each visual area. Principal component analysis (PCA) was used to reduce the dimension of the features while keeping 99% of the variance in each layer. We also built an encoding model with features from all CNN layers concatenated together. Again, PCA was used to keep 99% of the variance across all layers.

Formally, we model voxel *ν*’s response, *y*_*ν*_, as a linear weighted combination of the features *ϕ*^*l*^ from layer *l* of the CNN:

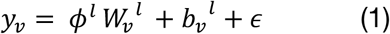

where *W*_*ν*_^*l*^ is the regression coefficient vector, *b*_*ν*_^*l*^ is the bias factor and ϵ is the model error. We then used Ridge regression with the following cost function to approximate regression coefficients from the training data:

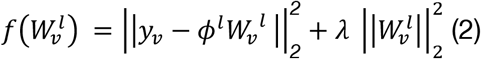

The regularization parameter *λ* is optimized through 10-fold cross-validation. After determining *λ*, the training data is used to estimate the final regression coefficients. Then, the prediction accuracy is obtained from the test set by calculating the average Pearson correlation coefficient between the predicted response and the measured response across test set segments.

### 2.3. Layer Contribution

For the encoding models from the entire CNN feature, we investigated the contribution of each CNN layer in predicting voxelwise responses. While it is possible to directly use the regression weights as the contribution, the values of the weights are highly dependent on the raw values of the feature sets [16]. Therefore, we calculated CNN layer contribution for each visual area as follows:

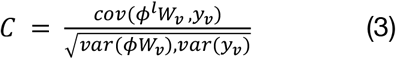

where *ϕ*^*l*^ is the feature map extracted from CNN layer *l, W*_*ν*_ is the regression coefficient vector, and 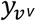 is the measured response.

### 2.4. Structural Compression

The process of structural compression involves removing redundant filters from the model to increase model interpretability. Here, we used a recently established structural compression technique called classification accuracy reduction (CAR) compression [25]. CAR compression quantifies the contribution of each filter to the model’s prediction accuracy and then removes the filters with the least contribution. We iteratively used CAR compression to continuously score and prune convolutional filters in each layer of AlexNet. Model accuracy was used as a stopping criterion to constrain the iterative structural compression i.e., the compression stopped when the hold-out validation set accuracy dropped 2% from the uncompressed model accuracy. Note that the hold-out validation set used for compression is different from the hold-out test set used for assessing the final accuracy. The compressed CNN-based features are then further compressed using PCA to retain 99% of their variance. These dimensionality-reduced features are then convolved with a canonical hemodynamic response function (HRF) [36] with the maximum at 5s and the outputs downsampled to match the fMRI sampling [8]. Finally, estimated fMRI responses are calculated with a ridge regression on the PCA-reduced, down-sampled features. At this step, the pruned model’s accuracy was determined by the Pearson correlation between measured and predicted responses on the hold-out test set. **Algorithm 1** summarizes the pseudo-code of the structural compression encoding model.

#### Algorithm 1 Structurally-compressed encoding model

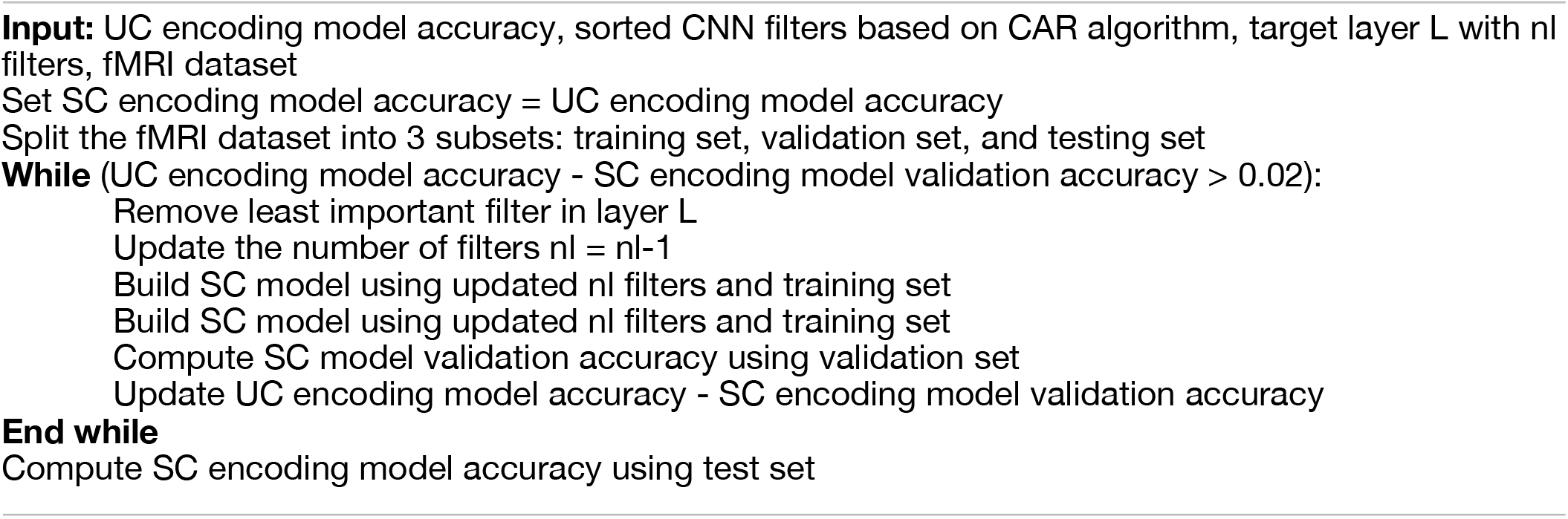

### 2.5. Deep Compression

We performed deep compression (DC) [27] to reduce the number of connections in the CNN-based encoding models. Following the process proposed in [27], we pruned the small-weight connections from the CNN. More specifically, all connections with weights below a threshold were removed while there was no drop in the classification accuracy. We further reduced the number of weights by having multiple connections share the same weight. We used k-means clustering to identify the shared weights in each layer of the network and used the same weights for all the connections that fell into the same cluster. We then fine-tuned the shared weights by retraining the network. Similar to structural compression, we further compressed the features using PCA followed by a convolution with the HRF. Finally, the BOLD responses were calculated using a ridge regression model. The model performance was assessed by the Pearson correlation coefficient between measured and predicted responses. **Algorithm 2** presents the pseudo-code of the deep compression encoding model.

#### Algorithm 2 Deep-compressed encoding model

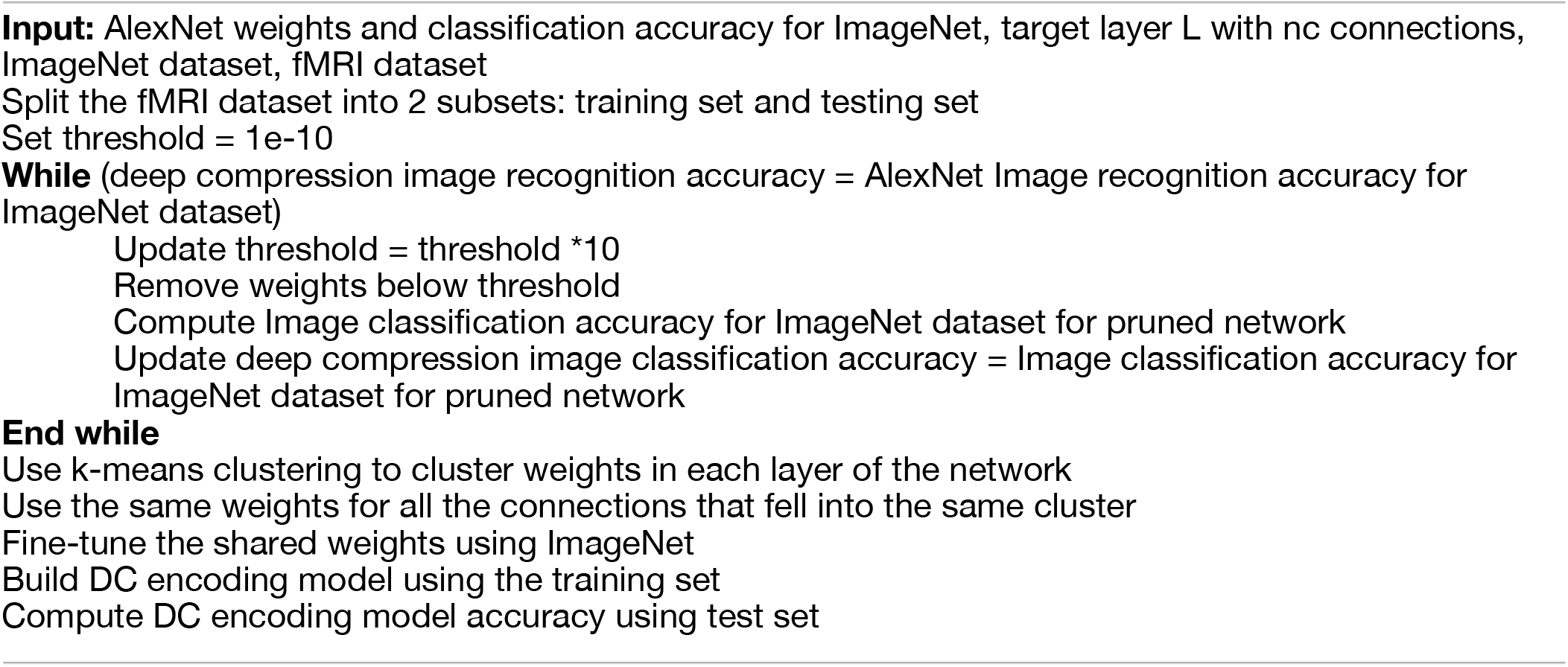

### 2.6. Receptive Field Compression

Receptive field compression takes inspiration from the biological visual pathways, much like how CNN architectures share similarities with the hierarchical organization of the visual cortex. This form of compression consists of identifying the most important regions of visual stimulus for the prediction task and then removing features from any location outside this region. Here, we modeled the population receptive fields with a 2D isotropic Gaussian function [16]. Thus, the population receptive field, *g*, can be described as:

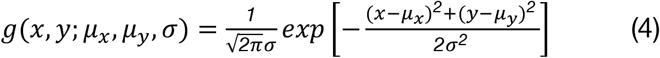

where (*μ*_*x*_, *μ*_*y*_) is the receptive field center and *σ* is the receptive field radius. To simulate the effect of biological receptive fields, the 2D Gaussian function was convolved with the CNN-based features extracted from the stimulus. Formally:

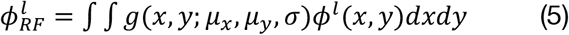

where 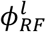 is the receptive field feature map for layer *l* and *ϕ*^*l*^ is the feature extracted from layer *l* of the CNN network.

We used grid search to approximate the optimal receptive field configuration for the CNN model. Since we’re using a 2D Gaussian function, we only varied the center and radius for the candidate receptive fields. For each voxel, we built a grid of candidate Gaussian pooling fields with varying sizes and locations. The size of the grid was 8 by 8 on the visual field (with Gaussian centers spaced 2.5 degrees apart). In each location on this grid, 8 log-spaced receptive fields were constructed with the sizes between *σ* = 0.5 and *σ* = 8. This provided a total of 512 Gaussian pooling fields on the visual field.

Once again, the Pearson correlation coefficient between measured and predicted responses was used to quantitatively pinpoint the best receptive field configuration. More specifically, we first applied each candidate receptive field to the features extracted from the CNN model, using Eq. 4. This was followed by a convolution with the HRF filter and downsampling to match the temporal frequency of the BOLD signal (0.5Hz for the PURR dataset, and 1Hz for the Vim-2 dataset). Similar to the structural and deep compression, PCA was used to reduce the feature dimension while keeping 99% of the variance. We used 80% of the dataset to train a Ridge regression model that predicts the fMRI BOLD signal from the compressed feature set. The prediction accuracy of this model was then assessed on the remaining 20% of the data. We used this process to identify the most accurate receptive field for each voxel. To determine the final accuracy of the compressed model, we retrained the Ridge regression using 100% of the training dataset and reported the accuracy on the hold-out test set for each voxel.

## 3. Results

### 3.1. Compressed CNN-based encoding models accurately predict BOLD responses

While our goal is to improve the interpretability of the CNN-based voxelwise models, we must also maintain the prediction accuracy of compressed models. To quantify prediction accuracy, we computed the voxelwise Pearson correlation coefficient between the predicted and measured BOLD signal. We then reported the average correlation coefficient for the following compression techniques: (1) structural compression (SC) where redundant filters are removed from the CNN-base model, (2) deep compression (DC) where CNN model weights that are close to zero are removed from the model, and (3) receptive field compression (RC) where the optimal receptive field size and location is determined for each voxel. Principal component analysis (PCA) was integrated with all these compression techniques to reduce the dimensionality of CNN features (see **Methods**). We have systematically compared the prediction accuracy of these compressed models with the uncompressed CNN-based model for each visual area. The visual areas that we considered are V1, V2, V3, V4, Lateral Occipital (LO), Middle Temporal (MT), Fusiform Face Area (FFA), Parahippocampal Place Area (PPA), Extrastriate Body Area (EBA), and Retrosplenial Cortex (RSC).

On average, the structurally- and deep-compressed models performed as well as the uncompressed model for most visual areas. The accuracies for the best-performing models among 8 distinct models that are built based on each layer of the CNN are presented in **Figures 1B and 1C**, while the accuracies using the entire set of CNN features from all layers are presented in **Figures 1D and 1E**. The average correlation coefficient is reported across all the voxels within each visual area. For FFA, the deep-compressed model underperforms other models by 3%, most likely due to the higher sensitivity of face features to the pruning of CNN weights. The receptive field compressed model has a slightly lower accuracy in the early and intermediate visual areas compared to other models (10% in V1, 5% in V2 and V3, and 2% in V4), but achieves similar accuracy to the uncompressed model for the higher visual areas. **Figure 1C** illustrates the predictive accuracy of the models based on each individual CNN layer for structurally-compressed models (as opposed to **1B** where the best-performing model is selected). The result is similar for other compression techniques (data not shown to avoid redundancy). Our findings suggest that early to middle layers of the CNN are better predictors of the responses in early and intermediate visual areas, while responses in higher visual areas are better predicted by deeper CNN layers. For areas V1, V2, V3, and V4, CNN layers 2 to 5 achieve the highest accuracies. For areas, LO, MT, FFA, PPA, and EBA, models based on layers 6 to 8 are the most accurate.

We further used the features extracted from all CNN layers in one single encoding model ±to estimate the BOLD responses. In addition to considering each individual compression technique, here we also present two combined compression methods: (1) structural and receptive field compression (SRC) and deep and receptive field compression (DRC). For both SRC and DRC, receptive-field compression is used after the structural or deep compression. The prediction accuracies reported in **Figure 1D** suggest that using the entire feature map to predict the BOLD signal results in higher accuracies (92% on average) compared to regressing single layers of CNNs. This is not surprising because a larger number of features with complimentary image statistics (from different layers of CNN) are used in these models compared to the single-layer models.

For the best-performing models in **Figure 1D**, we also quantified the contribution of each layer to the prediction accuracy for each visual area (**Figure 1E**). Overall, our results suggest that the average contribution of the extracted feature maps to predicting the BOLD signal in each visual area largely depends on the position of that area in the visual hierarchy. This is consistent with observations from previous studies [13] indicating that features from lower CNN layers have a higher contribution to the prediction of responses in lower visual areas, while higher visual areas are better predicted by the higher CNN layers.

We further visualize the voxelwise accuracies based on the entire CNN layers on the cortical map in one of the subjects (see **Supplementary Materials, Figure A1-7** for the accuracies in each individual subject). To create these maps we computed the Pearson correlation coefficients between predicted and measured responses for each voxel. The inflated and flattened cortical maps for the uncompressed and compressed models are shown in **Figure 2**. The structurally-compressed model outperforms the uncompressed model in ProS, DVT, and the lateral part of the V1, suggesting that these regions are better modeled using fewer CNN filters. The deep-compressed model has a higher predictive accuracy compared to the uncompressed model in parts of V1, V2, V3, and the lateral part of V4. For the receptive field-compressed model, the lateral parts of V1, V2, V3, V4, and also TPOJ and FST areas are modeled more accurately compared to the uncompressed model. Other subjects display similar results (**Supplementary Materials Figure A1-7**).

**Figure 2:**
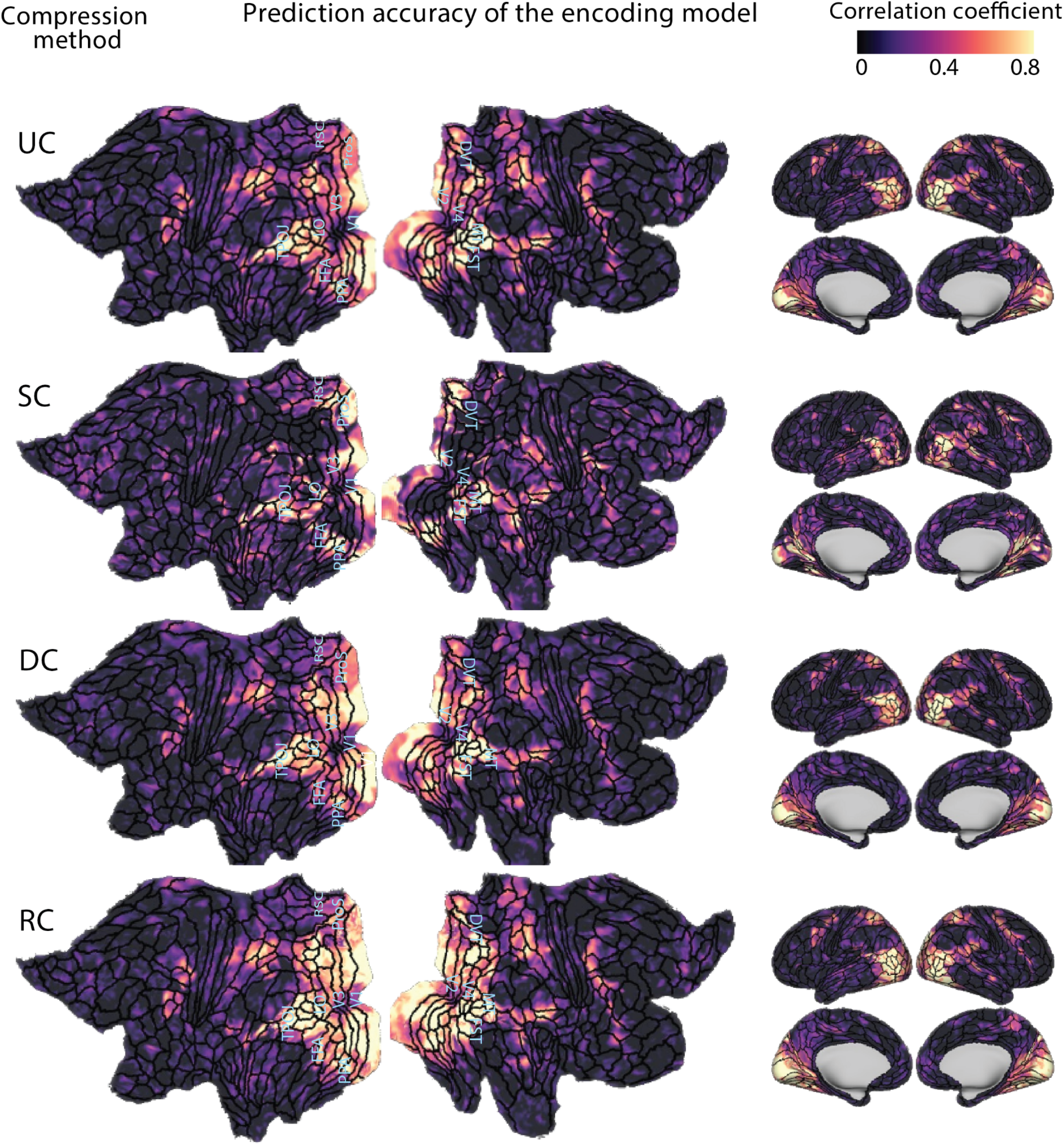
Prediction accuracy of the compressed encoding models on cortical maps. The maps show the correlations between estimated and measured responses for the uncompressed model (UC), structurally-compressed model (SC), deep-compressed model (DC), and receptive field-compressed model (RC) for subject 1 in the PURR dataset. Compared to uncompressed models, structurally-compressed models better estimate fMRI responses in ProS, DVT, and the lateral part of V1. The deep-compressed model is better in the central part of V1, V2, and V3, and the lateral part of V4. The receptive field compressed model better estimates the lateral part of V1, V2, V3, and V4, as well as TPOJ and FST areas.

### 3.2. Compression reduces the model size and computational cost while preserving the accuracy

So far, we have demonstrated that the compressed encoding models (see **Methods**) retain a satisfactory prediction accuracy. Here, we examine the model size, the computational cost, and the compression ratio of these compressed models. The model size is quantified by the number of weights (or parameters) in the model. To quantify the computational cost, we tracked the number of floating point operations per second (FLOPS) and the number of trainable parameters. For a convolutional layer in a neural network, the number of FLOPS refers to the number of floating point operations in that layer to extract all of the feature maps, without accounting for the regression overhead. The compression ratio was computed by dividing the number of FLOPS (or weights) required for the uncompressed model by that of the compressed model.

Our compressed encoding models have a reduced model size and a remarkably lower computational cost compared to the uncompressed models (**Table 1**). For structural compression, the model based on the CNN layer 4 has the highest compression ratio (compression ratio of 2) and the model based on layer 5 has the lowest compression ratio (ratio of 1.2), suggesting a high redundancy across layer 4 filters. Note that the compression ratio for structural compression is reported for a compressed model with a validation set accuracy that is less than 2% of the uncompressed model (see Methods). For the deep compression, the model based on the second fully connected layer has the highest compression ratio in terms of the number of FLOPS (ratio of 17.12), while models based on both the first and second fully connected layers have the highest compression ratio in terms of the number of weights (ratio of 33.33). These extraordinarily high compression ratios suggest the considerable amount of weight redundancy in fully connected layers of CNN. The last row in Table 1 shows the FLOPS, number of trainable weights, and compression ratios for the model that uses features from all layers of the AlexNet. For this model, structural compression has a compression ratio of 1.8 while deep compression has a ratio of 2.4 (FLOPS) and 27 (weights). For this model, the high compression ratio for the weights is due to the inclusion of fully connected layers in the model design. The PCA and the receptive field compression methods are not included in this table because these methods only compress the regression module, therefore the number of FLOPS does not change compared to the uncompressed models. However, it is worth noting that across different compression techniques, PCA explains 99% of the variance using 6664 to 10917 PCs for the convolutional layers, 2969 to 3463 PCs for the fully connected layers, and 241 components for the final softmax layer. For the models that are built based on all CNN layers, PCA explains 99% of variance using 1320 -2716 PCs across different compression techniques. Overall, our findings suggest that compression methods offer a reduced model size and computational cost.

**Table 1:** Comparison of the computational cost and the number of weights for the uncompressed, structurally-compressed, and deep-compressed models. The computational cost is quantified by FLOPS, i.e. the number of floating-point operations required in each layer to classify one image. The compression ratio is defined as the number of FLOPS (or weights) required for the uncompressed model divided by those required for the compressed model. Note that for the structurally-compression model, the compression ratio based on the number of FLOPS is equal to the compression ratio based on the number weights because all filters are removed during this form of compression. Structural compression is also not defined for the fully connected layers; therefore, no number is reported for these layers.

| Layer | Compression method | #FLOPS | Compression ratio (FLOPS) | #Weights | Compression ratio (Weights) |
| --- | --- | --- | --- | --- | --- |
| conv1 | Uncompressed | 105.41M | - | 35K | - |
|  | Structural | 58.56M | 1.8 | 19.4K | 1.8 |
|  | Deep | 88.89M | 1.18 | 11.43K | 3.06 |
| conv2 | Uncompressed | 223.95M | - | 307K | - |
|  | Structural | 124.41M | 1.8 | 170.55K | 1.8 |
|  | Deep | 86.38M | 2.59 | 44.55K | 6.89 |
| conv3 | Uncompressed | 149.52M | - | 885K | - |
|  | Structural | 93.45M | 1.6 | 553.12K | 1.6 |
|  | Deep | 52.24M | 2.86 | 115.98K | 7.63 |
| conv4 | Uncompressed | 112.14M | - | 663K | - |
|  | Structural | 56.07M | 2 | 331.5K | 2 |
|  | Deep | 41.89M | 2.67 | 93.51K | 7.09 |
| conv5 | Uncompressed | 74.76M | - | 442K | - |
|  | Structural | 62.3M | 1.2 | 368.33K | 1.2 |
|  | Deep | 27.69M | 2.69 | 61.09K | 7.14 |
| fc6 | Uncompressed | 37.75M | - | 38M | - |
|  | Deep | 2.33M | 16.18 | 1.14M | 33.33 |
| fc7 | Uncompressed | 16.78M | - | 17M | - |
|  | Deep | 0.98M | 17.12 | 0.51M | 33.33 |
| fc8 | Uncompressed | 4.10M | - | 4M | - |
|  | Deep | 0.53M | 7.71 | 0.29M | 13.69 |
| All layers together | Uncompressed | 724.37M | - | 61M | - |
|  | Structural | 394.79M | 1.8 | 33.8M | 1.8 |
|  | Deep | 300.79M | 2.4 | 2.25M | 27 |

### 3.3. Compressed models identify natural movie frames from BOLD responses more accurately than uncompressed models

Having established that compressed models reduce computational cost while retaining a high predictive accuracy compared to their uncompressed counterparts, we now examine how compression may improve the interpretability of voxelwise encoding models. More specifically, we investigate whether compression improves the model’s ability to decode visual stimuli from brain activity (BOLD signal). To test this, we compared the ability of the compressed and uncompressed models to identify which video frame was shown to the subject during a hold-out test set with 598 clips. We first used structurally-compressed (SC), deep-compressed (DC), receptive field-compressed (RC), and the uncompressed (UC) model to predict voxelwise activity patterns evoked by each clip in the hold-out test set. We then selected the video clip with the highest Pearson correlation coefficient between predicted and the measured BOLD signal (**Fig. 3**). The identification performance was obtained as the ratio of the number of correctly selected clips to the total number of clips. The average identification performance across all subjects in the PURR dataset is 91%, 90.6%, 92%, and 93% for UC, SC, DC, and RC models, respectively. For the vim-2 dataset, the identification performance was 81.6%, 83.3%, 85%, and 87% for UC, SC, DC, and RC models, respectively. Overall, the RC model performed better in identification accuracy than the uncompressed model and the other compressed models. Note that the chance-level performance is 0.1% for subjects in the PURR dataset and 1% for the subjects in the vim-2 dataset.

**Figure 3:**
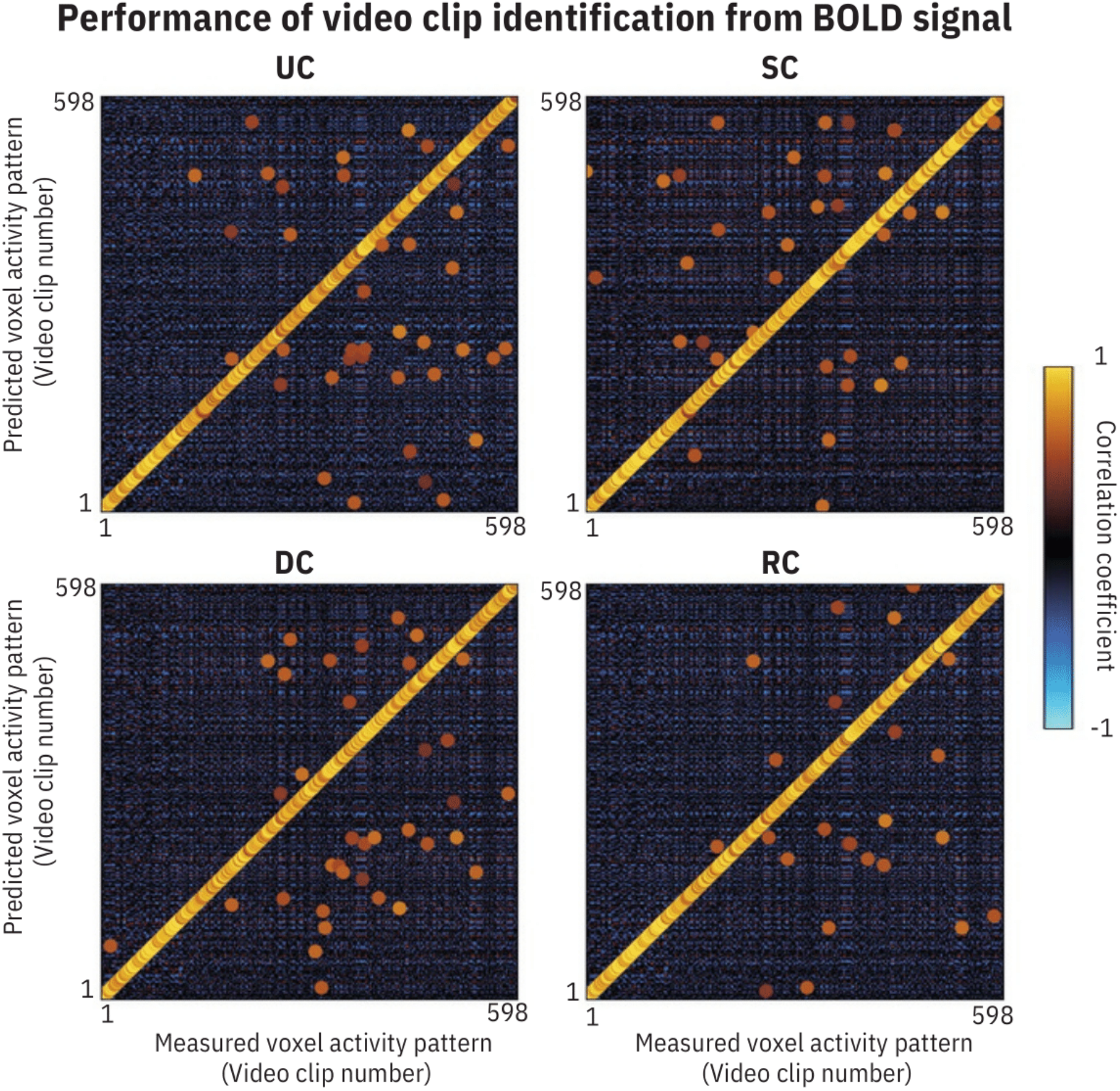
Performance of video clip identification from BOLD signal. The Pearson correlation coefficient between the measured BOLD signal and the predicted BOLD signal for subject 1 in the PURR dataset. The structurally-compressed (SC), deep-compressed (DC), receptive field-compressed (RC), and uncompressed (UC) models were used to predict the BOLD signal. In each column, the highest correlation coefficient is identified with an enlarged circle and is colored according to its associated correlation coefficient. A diagonal pattern suggests high identification performance.

### 3.4 Structurally-compressed models reveal more stable interpretations compared to the uncompressed models

Another established method for the interpretability of a voxelwise model is visualizing the images that elicit the largest response in each voxel (i.e., optimal visual patterns) [3]. This form of interpretability may be hindered for CNN-based models due to their large number of parameters [25]. More specifically, the optimal visual patterns for each voxel could be *unstable* when a large model is used to identify or generate that pattern [37]. Here, stability is defined as the similarity of visual patterns among the images with the highest model responses. This form of stability is a necessity for the reliable interpretation of the CNN-based models for scientific discovery [22]. Considering the lower number of parameters in the structurally-compressed model, we hypothesize that the images that elicit the largest response in each voxel are more stable for the structurally-compressed model compared to the uncompressed model.

To determine the optimal visual pattern for each voxel, we searched for the natural images that elicit the largest response for both the uncompressed and structurally-compressed models. We chose 10,000 natural images from the validation set in the ILSVRC 2012 dataset [34] and computed the voxelwise response of both encoding models to each image. The images with the highest response were selected for each voxel. Note that these images were neither used to train the CNN nor included in the experimental stimulus set. **Figure 4A** presents the top 5 images with the highest model response for the most accurately predicted voxel in each visual area. We have visualized these preferred images for the top 10 most accurately predicted voxels in each visual area in **Supplementary Materials, Figures A8-12**. Overall, these images provide a qualitative representation of the patterns selected by each voxel. The patterns for the voxels in V1 and V2 do not contain category-specific patterns. This is consistent with previous observations that V1 and V2 areas lack category selectivity and are more selective to lower-level image features such as edges and borders [3, 13, 38]. The top images for the V4 area contain semi-complex shapes such as curvatures, circles, crosses, and dense textures. For FFA, the top 5 images include face features while the top images for PPA and RSC contain environmental scenes. These findings are aligned with observations in previous studies [37], suggesting both uncompressed and compressed models are successful in determining the preferred images for the accurately predicted voxels.

**Figure 4:**
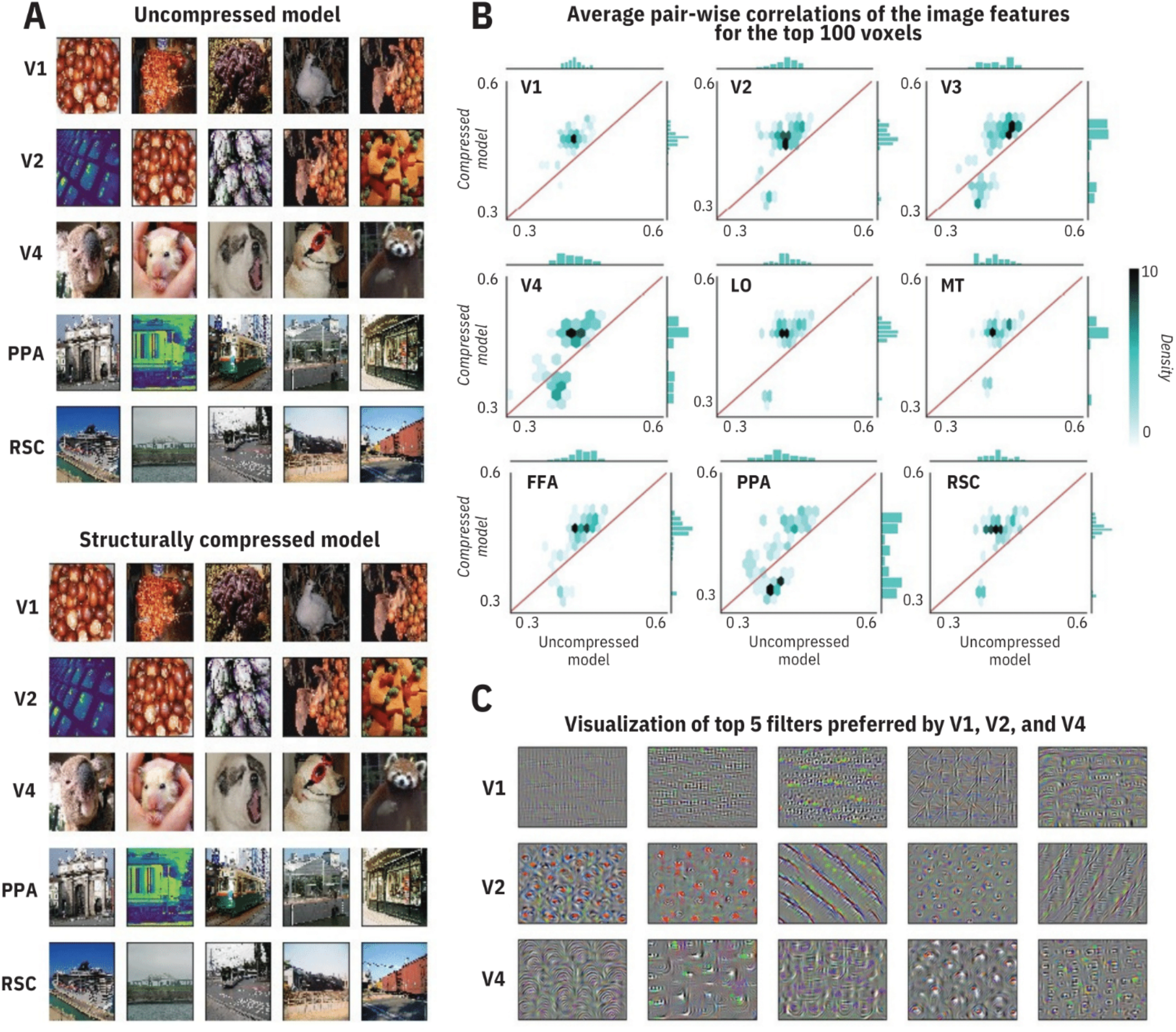
Structurally-compressed models reveal more stable category selectivity compared to the uncompressed models. **A**. Top 5 images with the highest model response in each visual area for subject 1 in the PURR dataset. Top images for the V1 and V2 areas are rich in lower-level image features such as edges and borders. The top images for the V4 area contain semi-complex shapes such as curvatures, circles, crosses, and dense textures. For FFA, the top five images include face features. The top images for PPA and RSC contain environmental scenes. **B**. Density plots comparing the stability of the selected images between compressed and uncompressed models. The stability is quantified by the average pairwise correlation coefficient between CNN-extracted features of the top 10 selected images for each voxel. The average correlation coefficient is reported across the top 100 voxels for 9 visual areas. The color of each hexagonal bin indicates the density of voxels in that bin. The histograms below each plot represent the distribution of the relative stability score. Overall, top images are more stable for the structurally-compressed encoding model compared to the uncompressed model. **C**. Visualization of the top filters for top voxels in the early ventral visual pathway. To visualize each filter, we performed a gradient ascent to create an image that maximizes the activation of that filter. Overall, V1 and V2 are tuned to edges in various directions while V4 is primarily tuned to curvatures. The results are from subject 1 in the PURR dataset.

We then examined the stability of the preferred stimulus images for each voxel and compared the stability between the compressed and uncompressed models. We used the similarity between CNN-extracted features from images to quantify the stability. This feature-based similarity measure is chosen over the pixel-based measure because CNN-extracted features encompass the overall content of each image and reflect the image patterns more reliably compared to the pixel values.

Here, we constructed the feature space by concatenating features from all layers of the AlexNet [34]. Formally, the stability score is defined as the average pairwise Pearson correlation coefficient between CNN-extracted features across images. We considered the stability score computed over the top 10 images with the highest model response. We computed this score for the top 100 most accurately predicted voxels in each area for both structurally-compressed and uncompressed models.

Scatterplots comparing these stability scores indicate that the stability is considerably higher for the structurally-compressed model in V1, V2, V3, V4, LO, MT, FFA, and RSC outperforming the uncompressed model by 0.07, 0.06, 0.04, 0.05, 0.06, 0.04, 0.05, and 0.06, respectively (**Figure 4B**). The stability scores in PPA are only marginally higher for the compressed model, outperforming the uncompressed model by only 0.008. Considering only the top 50 most accurately predicted voxels in PPA, the compressed model dominates the uncompressed model – the compressed model has a higher correlation coefficient for more than 81% of the top voxels across all visual areas. These results indicate that the compressed model provides more stable features than the uncompressed model for the majority of accurately predicted voxels.

We then asked whether the stability is a consequence of structural compression or any reduction in the number of parameters. To answer this question, we repeated the stability analysis for randomly-pruned filters with the same compression rate as structural compression. Averaged over 100 repeats of this random pruning procedure, the stability score dropped by 8% for V1, 8% for V2, 6% for V3, and 4% for V4, suggesting the significance of structural compression in improved stability for these areas. The stability score remained the same for RSC, LO, MT, FFA, and PPA, which is not surprising because these areas are better modeled by fully connected layers of CNN (layers 6, 7, and 8, see **Figure 1B**) which are not affected by structural compression.

To further investigate the preferred visual patterns in each visual area, we synthesized the optimal patterns for the top CNN filters preferred by each visual area (**Figure 4C**). First, we selected the top five CNN filters for each visual area by identifying the filters with the highest predictive performance for all the voxels in that visual area. For example, for V1, the best-performing model uses the third convolution layer in AlexNet with 384 filters. We made 384 encoding models using each of these filters separately and calculated the average correlation coefficient between predicted and measured responses for all voxels in V1. The top 5 filters with the highest accuracies were selected as filters preferred by V1. We then synthesized the optimal visual pattern for each of these filters by direct optimization of the input stimulus image to maximize the filter response. The cost function in this optimization problem was defined as the mean activation of a target CNN filter. Starting from a random input image, we then used gradient ascent to generate the optimal image with the highest CNN filter response (with a learning rate of 1 for 1000 epochs). **Figure 4C** illustrates the visualization of these 5 synthesized patterns for the areas V1, V2, and V4. The accuracies for single-filter encoding models were 49%, 48%, and 35% for V1, V2, and V4, respectively. Optimal patterns for the areas V1 and V2 are diverse and include edges in various directions, while V4 tends to be tuned to curvature patterns. These observations are consistent with findings from **Figure 4A**.

Overall, our findings indicate that structurally-compressed models allow for a more stable interpretation of pattern selectivity for each voxel. Consistent with prior studies [3, 19], our findings indicate that the downstream areas in the ventral visual pathway are more category-selective compared to early areas.

### 3.5. The receptive-field-compressed models reveal increased size and centralization of the population receptive fields along the ventral visual pathway

The organization of the population receptive field maps (characterized by neural population-level measurements such as BOLD signal) in different areas of the visual cortex have been extensively studied in the past [39-40], [16]. These studies have provided evidence for larger and more centralized receptive fields in higher visual areas along the ventral visual pathway. We confirmed this observation using the receptive-field-compressed encoding models. These models allow for a systematic and quantitative analysis of the optimal size and location of the model-based population receptive fields in different visual areas. **Figure 5A** illustrates the selected population receptive fields for the top 100 most accurate voxels in visual areas V1, V2, V3, V4, LO, MT, FFA, PPA, EBA, and RSC. Visually, the areas in the early visual pathway have smaller and more scattered population receptive fields while higher visual areas have larger and more centralized receptive fields. To quantitatively compare the population receptive field locations and sizes, we computed the mean absolute distance between the center of each receptive field and the mean receptive field center across all voxels in each visual area (**Figure 5B**). We found that V1 has the smallest average population receptive field size (mean radius: 1.6±0.02 degrees) with the most scattered centers (mean eccentricity: 7.13±0.04 degrees). V2, V3, and V4 exhibit slightly larger mean receptive field size (mean radius: 1.75±0.02 and 1.77±0.02, and 1.86±0.03 degrees, respectively) with gradually increasing centralization (mean eccentricity: 7.08±0.04, 6.64±0.04, and 6.10±0.04 degrees, respectively). On the other hand, voxels in FFA and PPA had the largest population receptive fields (mean radius: 3.33±0.08 and 3.40±0.07 degrees, respectively) followed by LO, MT, and EBA (mean radius: 2.22±0.04 and 2.66±0.07, and 2.55±0.09 degrees, respectively). The population receptive fields in MT, FFA, PPA, and EBA were the most centralized (mean eccentricity: 5.27±0.09, 4.58±0.08, 4.68±0.07 and 4.78±0.09, respectively). Overall, the receptive field-compressed encoding model provides a quantitative and systematic framework to compare the population receptive field sizes and centers along the visual hierarchy. These models systematically confirm findings by previous studies that the population receptive fields become larger and more concentrated as we move downstream in ventral pathways.

**Figure 5:**
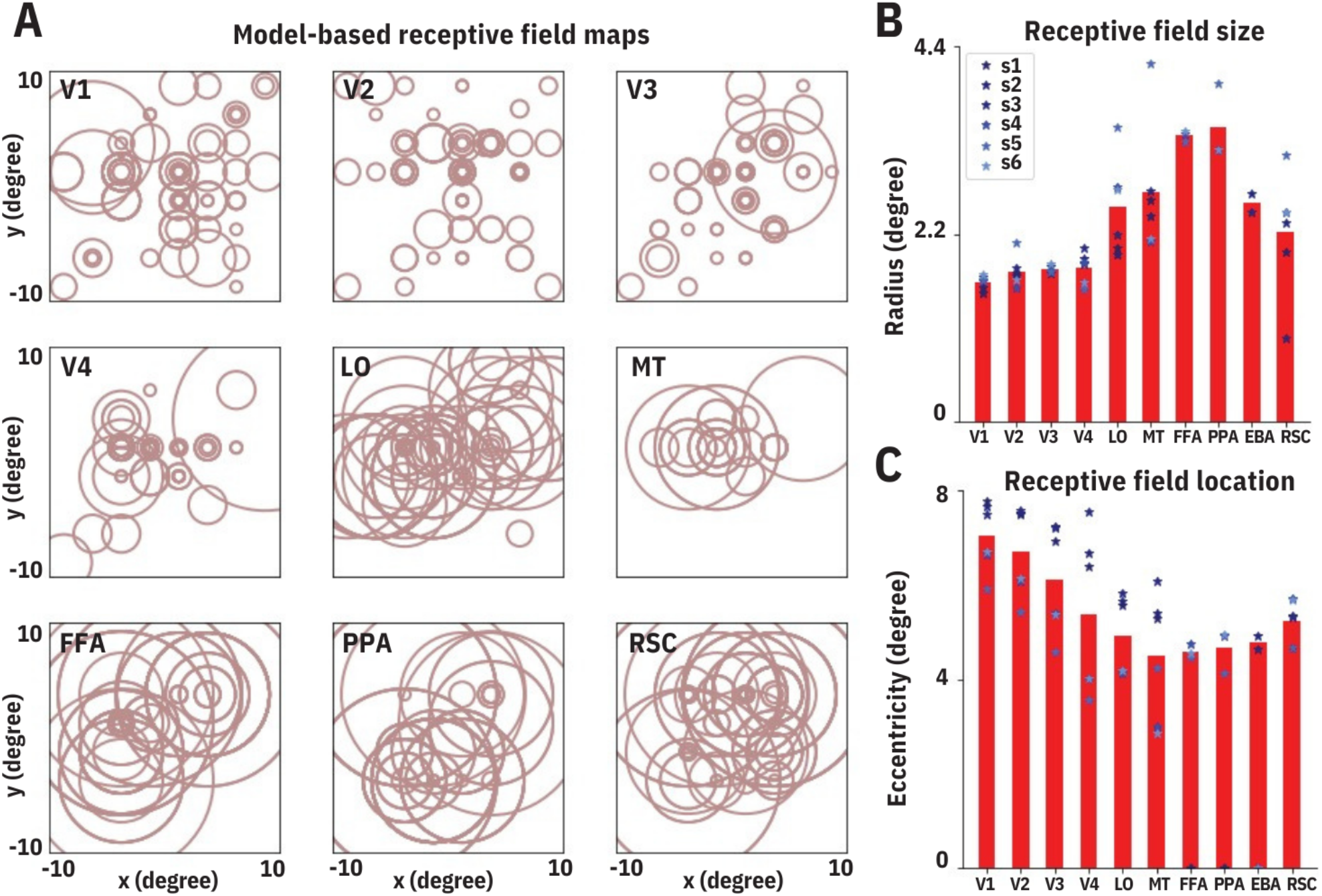
The receptive-field-compressed models reveal increased size and centralization of the population receptive fields along the ventral visual pathway. **A**. Each circle illustrates the receptive field for an individual voxel from the top 100 most accurately predicted voxels in subject 1 from the PURR dataset. The radius and the center of each circle are determined by the radius and the center of the Gaussian pooling field selected through receptive field compression. **B**. The mean radius of receptive fields in each visual area. **C**. The mean absolute distance between the center of each receptive field and the mean receptive field center across all voxels in each visual area. Receptive fields become larger and more concentrated as we move toward the downstream regions in the visual area.

## 4. Discussion

Encoding and decoding models are powerful tools to investigate human vision. We have shown that compressing CNN-based encoding models significantly decreases the number of parameters involved and reduces the computational cost while preserving accuracy. We used structural compression to remove less important filters, deep compression to remove less important connections, and receptive-field compression to pool the features. Our findings suggest that compressed encoding models provide an interpretable and quantitative framework to investigate the relationship between natural visual stimuli and the fMRI BOLD signal.

Overall, we demonstrated that the small set of visual stimulus features identified by compression could accurately predict the BOLD signal. However, modeling higher visual areas (e.g., FFA) that encode more complex visual patterns (e.g., faces) may still require a larger number of features. This is evident from our finding for FFA which showed high accuracy for a model that uses all layers of CNN (correlation coefficient of 0.52±0.009) compared to the best-performing individual-layer model (correlation coefficient of 0.30±0.009) (**Figure 1D**). This suggests that FFA models require a diverse set of features including both low-level and high-level image statistics, which is not surprising due to the complexity of the face images that are primarily encoded by FFA.

Our findings suggest that combining multiple compression strategies could increase the compression ratio while maintaining predictive accuracy (**Figure 1D**). Receptive field compression combined with structural compression, however, could introduce aggressive feature pruning in the design of the predictive model, leading to a partial reduction in predictive performance (**Figure 1D**). For the structural compression, our findings confirmed previous studies [25] that the convolutional layers in a CNN such as AlexNet contain redundant but diverse filters. Here we reduced the redundancy but maintained the diversity [25], which is key for the high accuracy of the structurally-compressed models, especially for early visual areas.

In terms of the encoding model design, we constructed a two-step encoding model that consists of an image stimulus feature extraction module and a BOLD response prediction module. The image feature extraction module was a deep convolutional neural network trained on a natural image classification task. An alternative modeling approach is to directly train a deep neural network that takes image stimuli as the input and predicts the voxelwise responses. This end-to-end model architecture could present a simpler model design and training but will require an order of magnitude larger fMRI dataset than what was used here. We expect that the compression methods used in this study would allow for training such an end-to-end model with limited fMRI data. This was beyond the scope of this study but is a direction for future follow-up studies.

Finally, the deep neural network used in this study was trained on a single task (image classification), however, the representations across the human visual cortex emerge in response to a variety of tasks such as classification, detection, and recognition. A more accurate encoding model would aggregate features relevant to these various tasks, but the huge size of the feature space in these aggregated models may introduce feasibility issues. Our findings suggest that future studies aimed at the construction of such multi-modal systems should consider compression techniques as an essential part of their design to decrease the feature space and allow for more effective model interpretation.

## Acknowledgments

The authors would like to thank Gavin Cui, Gaurav Ghosal, and the anonymous reviewers of the manuscript for their valuable feedback on the manuscript.

## Ethics Approval

Not applicable.

## Consent to Participate

Not applicable.

## Consent for Publication

Not applicable.

## Availability of Data and Materials

All the data used in this manuscript are publicly available at https://crcns.org/data-sets/vc/vim-2 and https://engineering.purdue.edu/libi/lab/Resource.html. The intermediate data files are available at tinyurl.com/nbds7cya.

## Code Availability

The software package is available at https://github.com/abbasilab/compressed-bold-models.

## Authors’ Contribution

R.A., A.S., and M.M. conceptualized and conceived the experiments. F.K. conducted the experiments. All authors analyzed the data. F.K. and R.A. wrote the manuscript with contributions from A.S. and M.M.

## A. Supplementary Materials

### A.1. Prediction accuracy of compressed encoding models on cortical maps for other subjects in the PURR dataset

**Figure A1** illustrates the voxelwise prediction accuracy on cortical maps for subjects 2 and 3 for the PURR dataset. The findings outlined in the main text for subject 1 are consistent with those for subjects 2 and 3.

**Figure A1:**
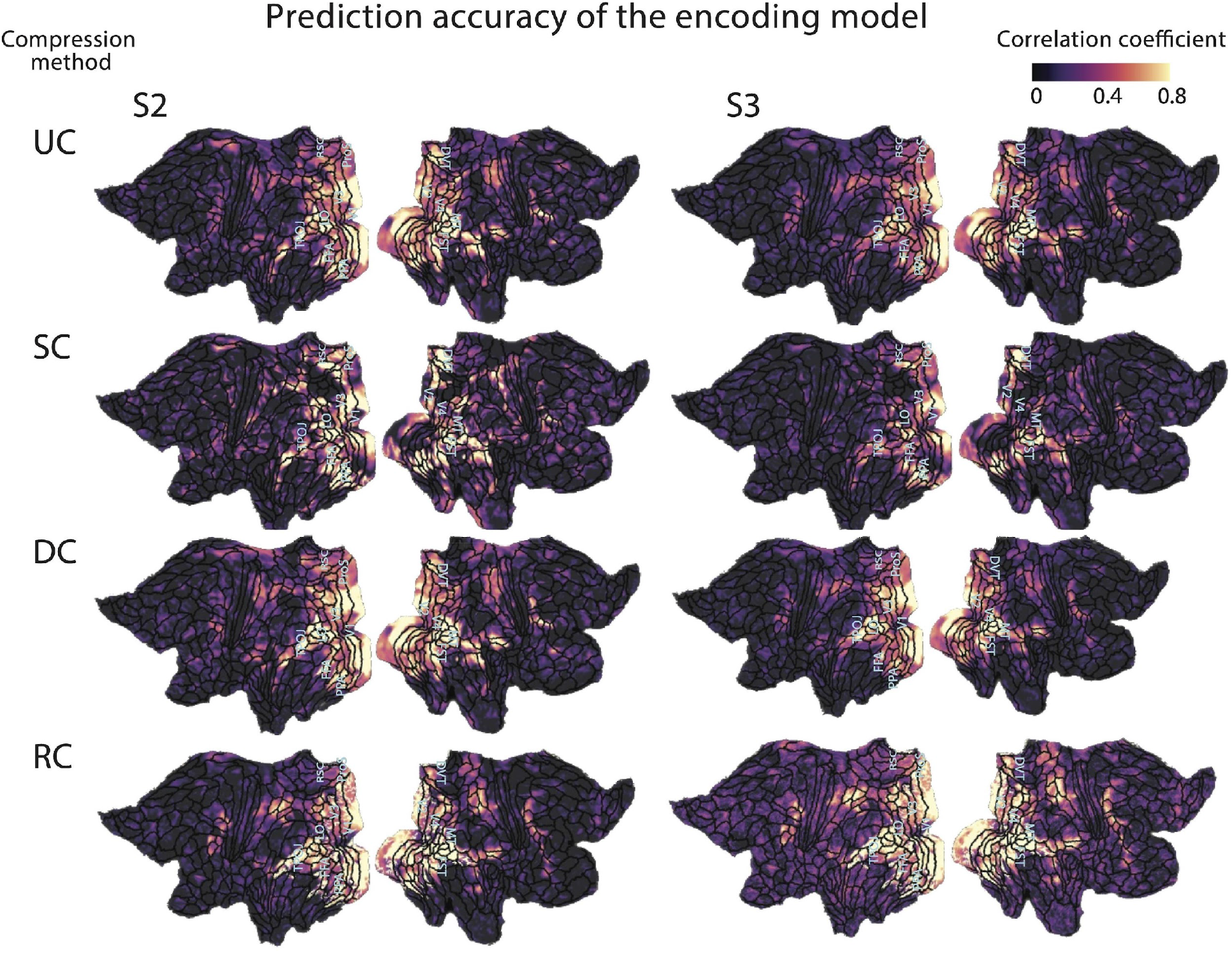
Prediction accuracy of compressed encoding models on cortical maps for subjects 2 and 3 in the PURR dataset. Compared to the uncompressed (UC) model, the structurally-compressed (SC) model better estimates fMRI responses in ProS, DVT, and the lateral part of V1. The deep-compressed (DC) model is more accurate in the central part of V1, V2, V3, and the lateral part of V4. The receptive field-compressed (RC) model better estimates the lateral part of V1, V2, V3, V4, as well as the TPOJ and FST areas.

### A.2. Voxelwise comparison of correlation coefficients between compressed and uncompressed models

**Figures A2-7** illustrate the voxelwise comparisons of correlation coefficients between the compressed and uncompressed models.

**Figure A2:**
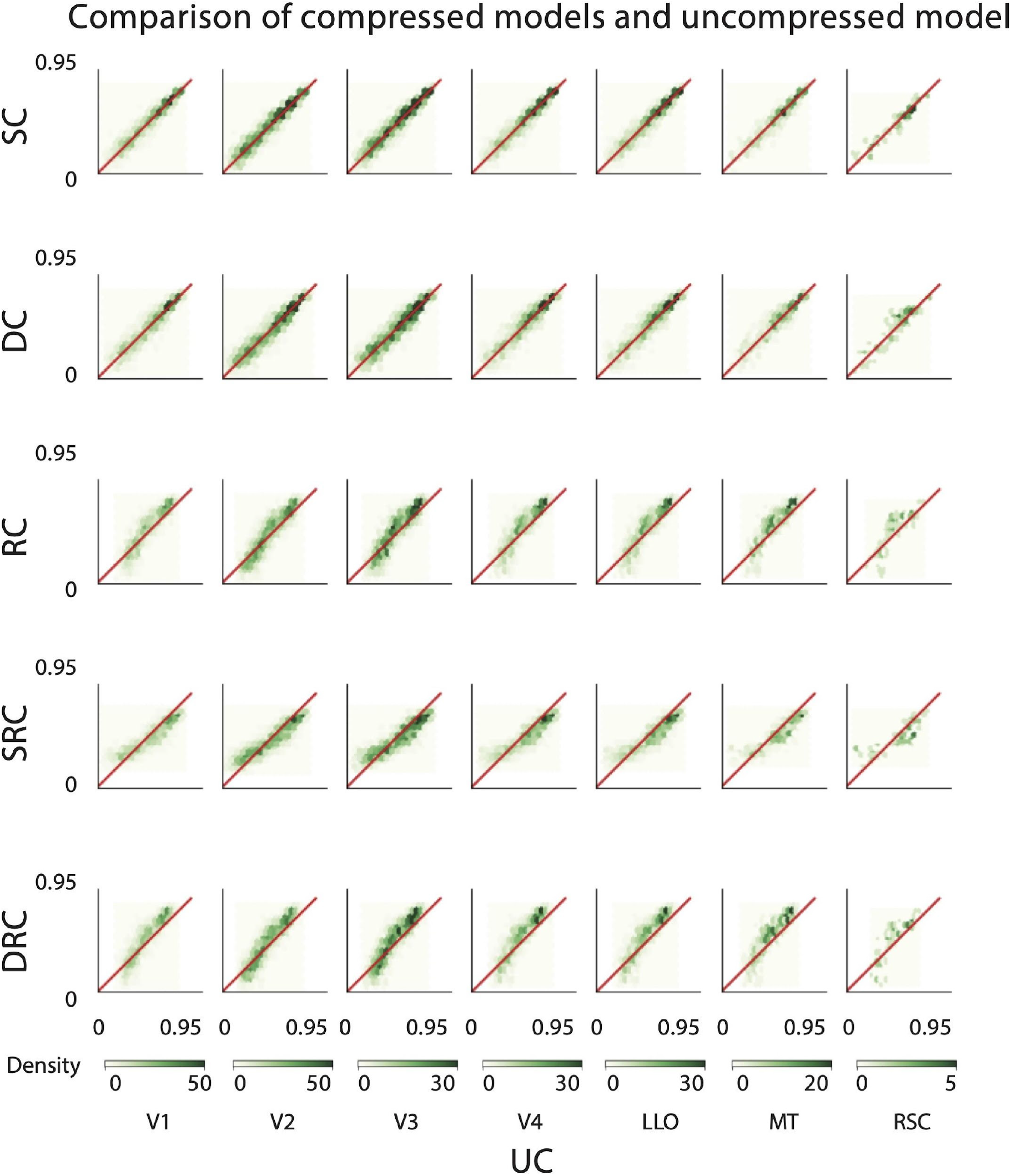
Scatterplots comparing voxel-wise correlation coefficients between compressed and uncompressed models for subject 1 in the Vim-2 dataset. Each dot corresponds to one voxel. Columns represent different visual areas. Rows represent different compression techniques.

**Figure A3:**
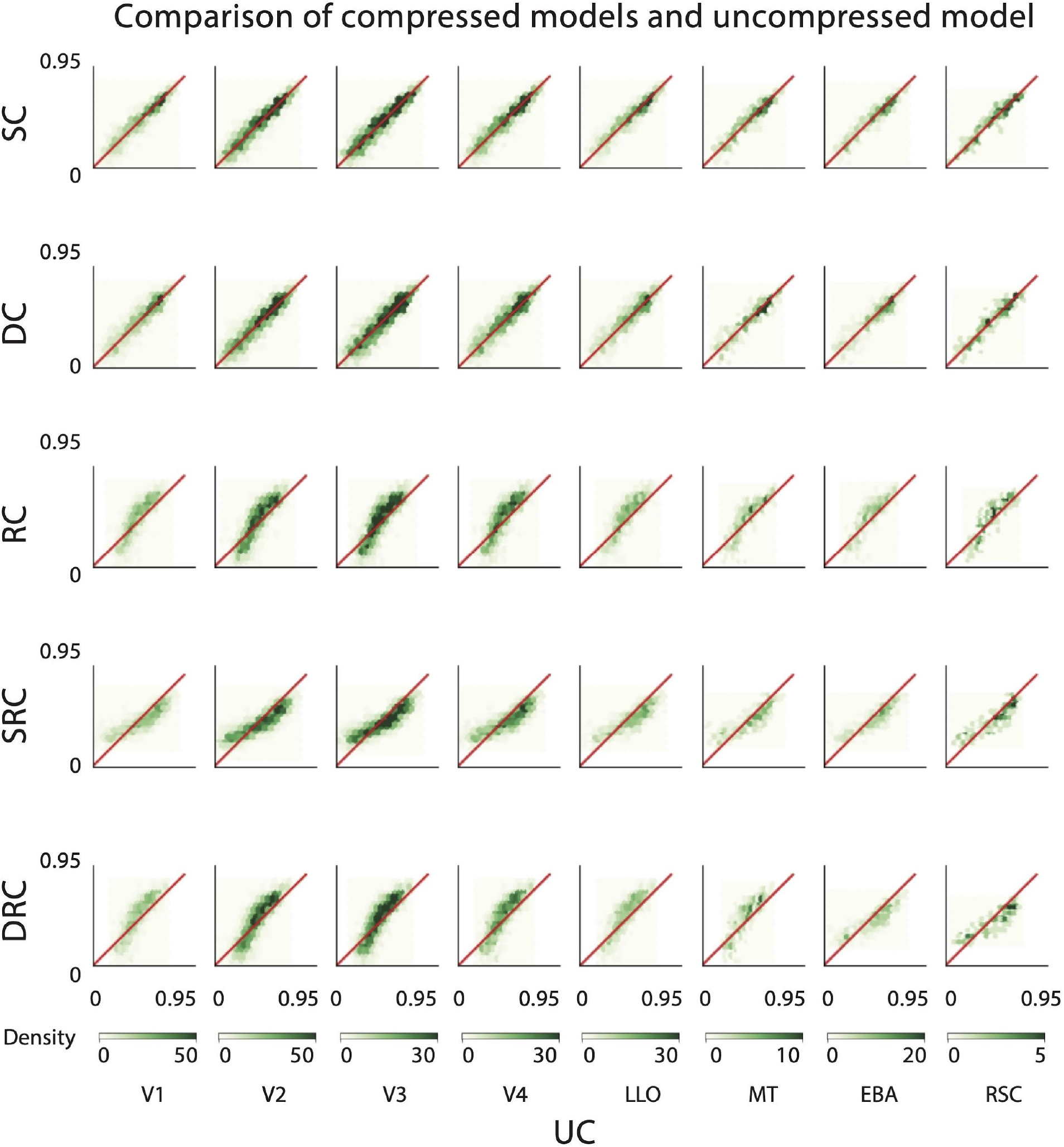
Scatterplots comparing voxelwise correlation coefficients between compressed and uncompressed models for subject 2 in the Vim-2 dataset. Each dot corresponds to one voxel. Columns represent different visual areas. Rows represent different compression techniques.

**Figure A4:**
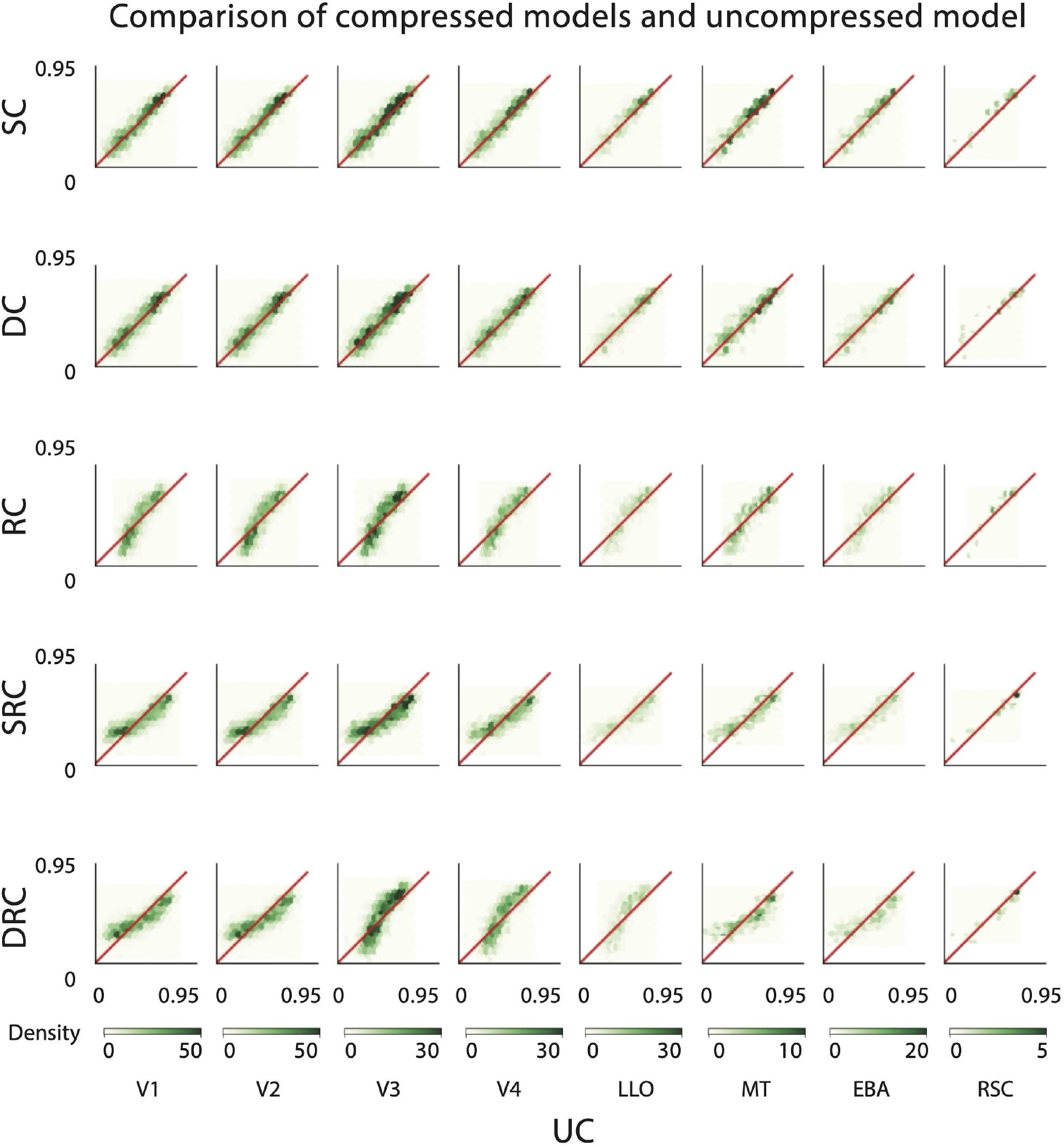
Scatterplots comparing voxelwise correlation coefficients between compressed and uncompressed models for subject 3 in the Vim-2 dataset. Each dot corresponds to one voxel. Columns represent different visual areas. Rows represent different compression techniques.

**Figure A5:**
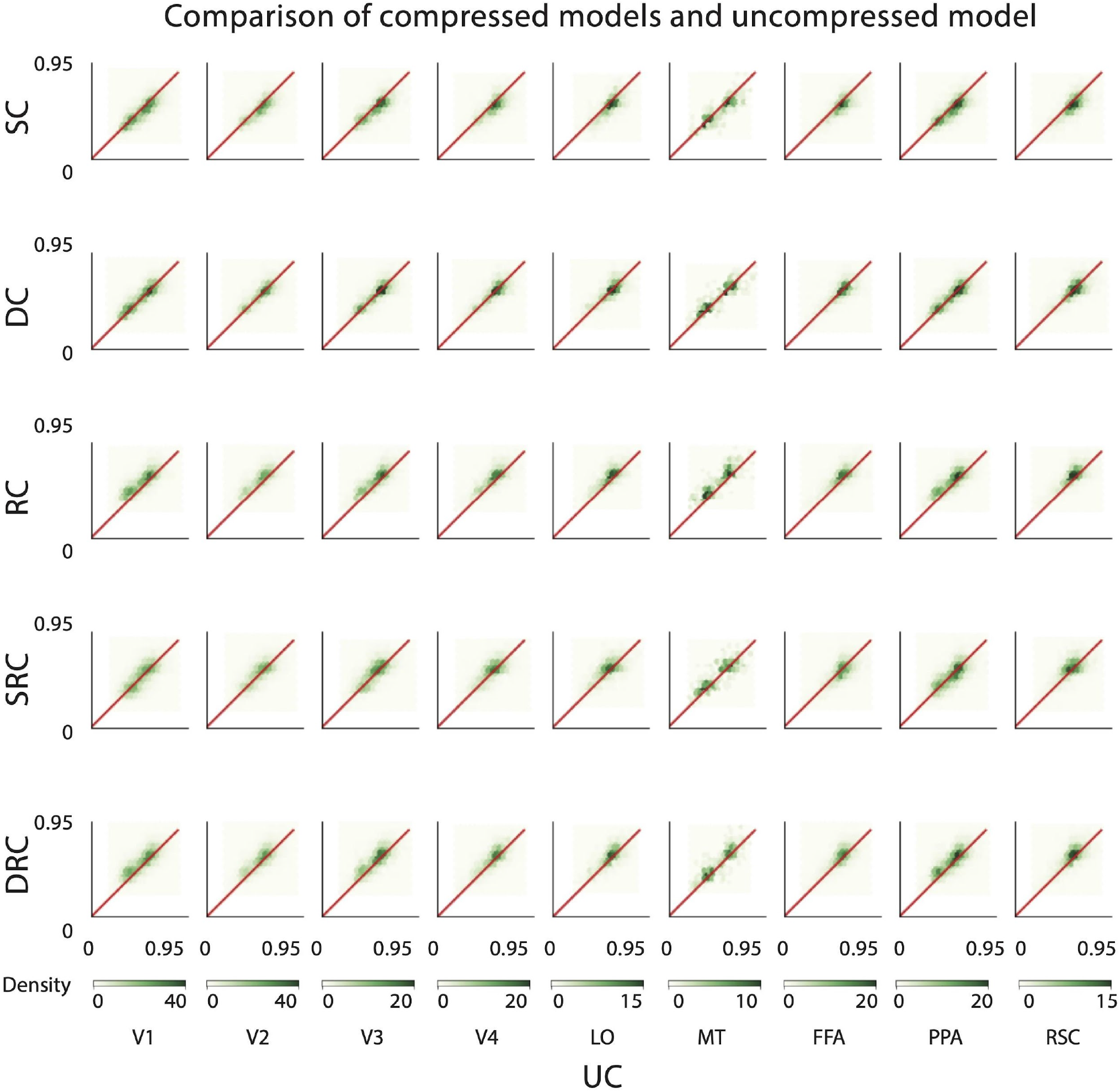
Scatterplots comparing voxelwise correlation coefficients between compressed and uncompressed models for subject 1 in the PURR dataset. Each dot corresponds to one voxel. Columns represent different visual areas. Rows represent different compression techniques.

**Figure A6:**
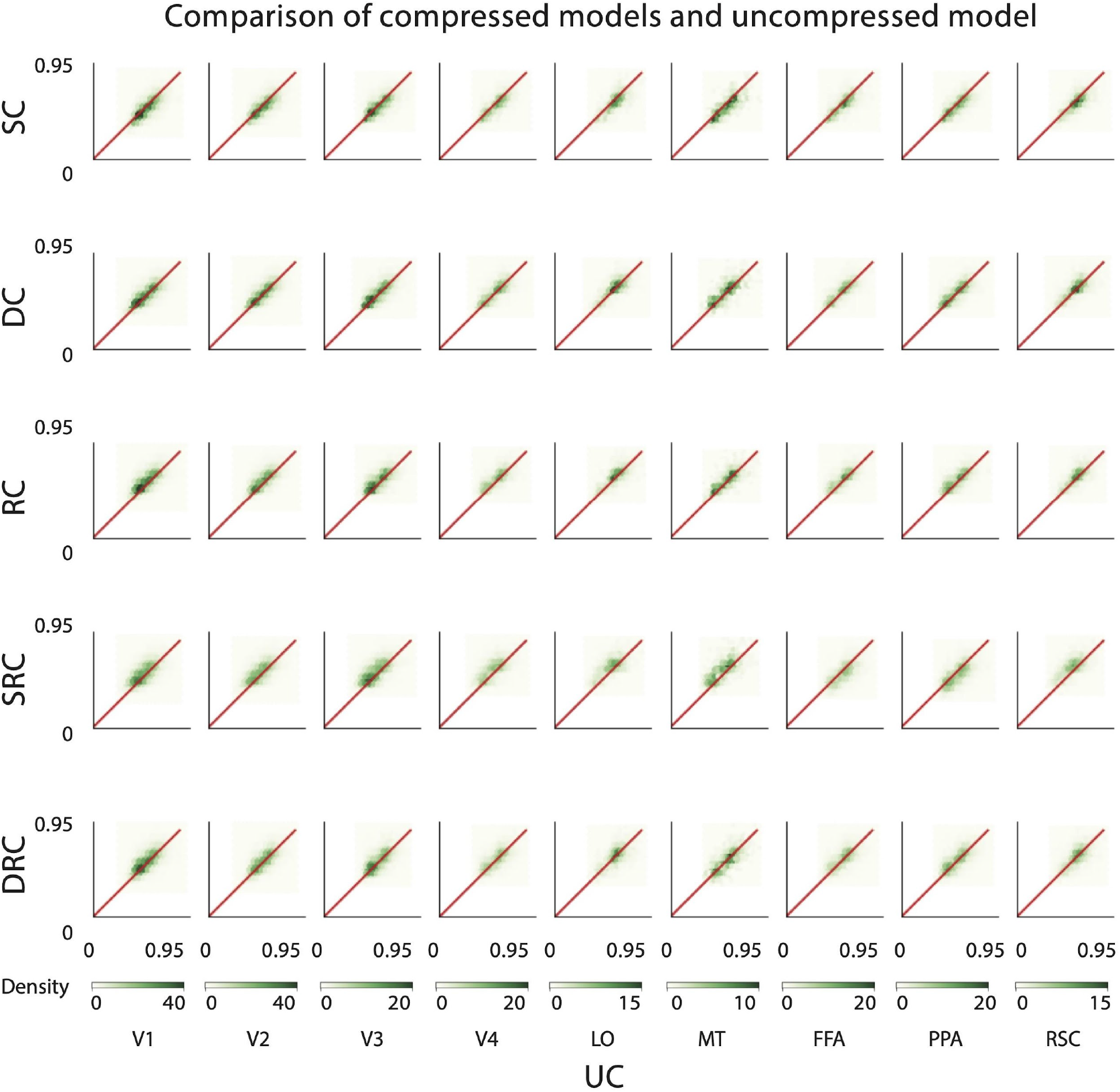
Scatterplots comparing voxelwise correlation coefficients between compressed and uncompressed models for subject 2 in the PURR dataset. Each dot corresponds to one voxel. Columns represent different visual areas. Rows represent different compression techniques.

**Figure A7:**
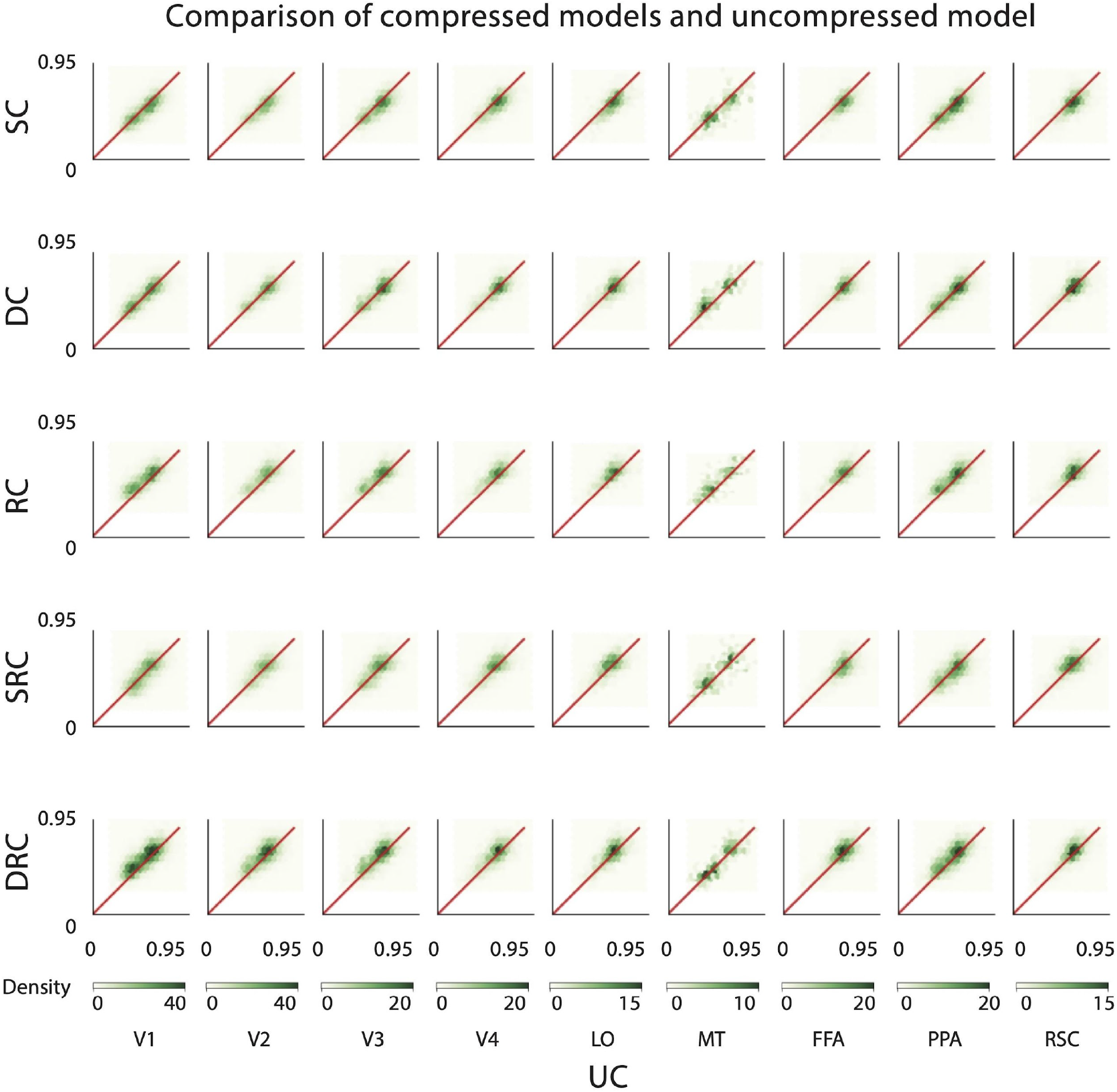
Scatterplots comparing voxelwise correlation coefficients between compressed and uncompressed models for subject 3 in the PURR dataset. Each dot corresponds to one voxel. Columns represent different visual areas. Rows represent different compression techniques.

### A.3. Top images with the highest model response for the structurally-compressed and uncompressed models

**Figures A8-13** present the 5 top images with the highest response for the top 10 most accurately voxels in visual areas V1, V2, V4, FFA, PPA, and RSC, respectively. Images are shown for both uncompressed and structurally-compressed models.

**Figure A8:**
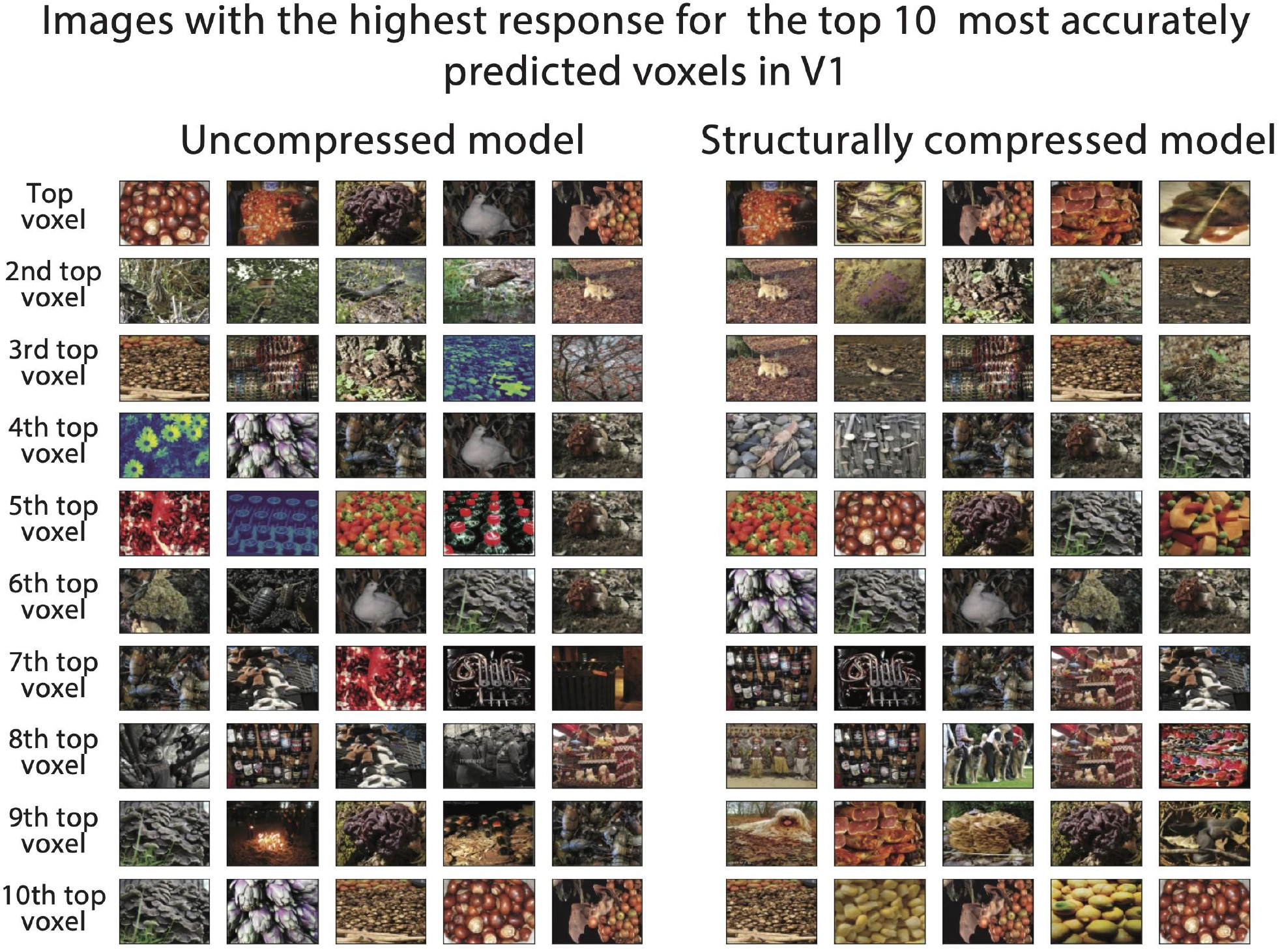
Comparing the performance of structurally-compressed and uncompressed models in V1. A. The five columns at left show the five images that the uncompressed model predicts will most increase activity for voxels in the V1 area, and the five columns at right show the five images that the structurally-compressed model predicts will most increase activity for voxels in the V1 area.

**Figure A9:**
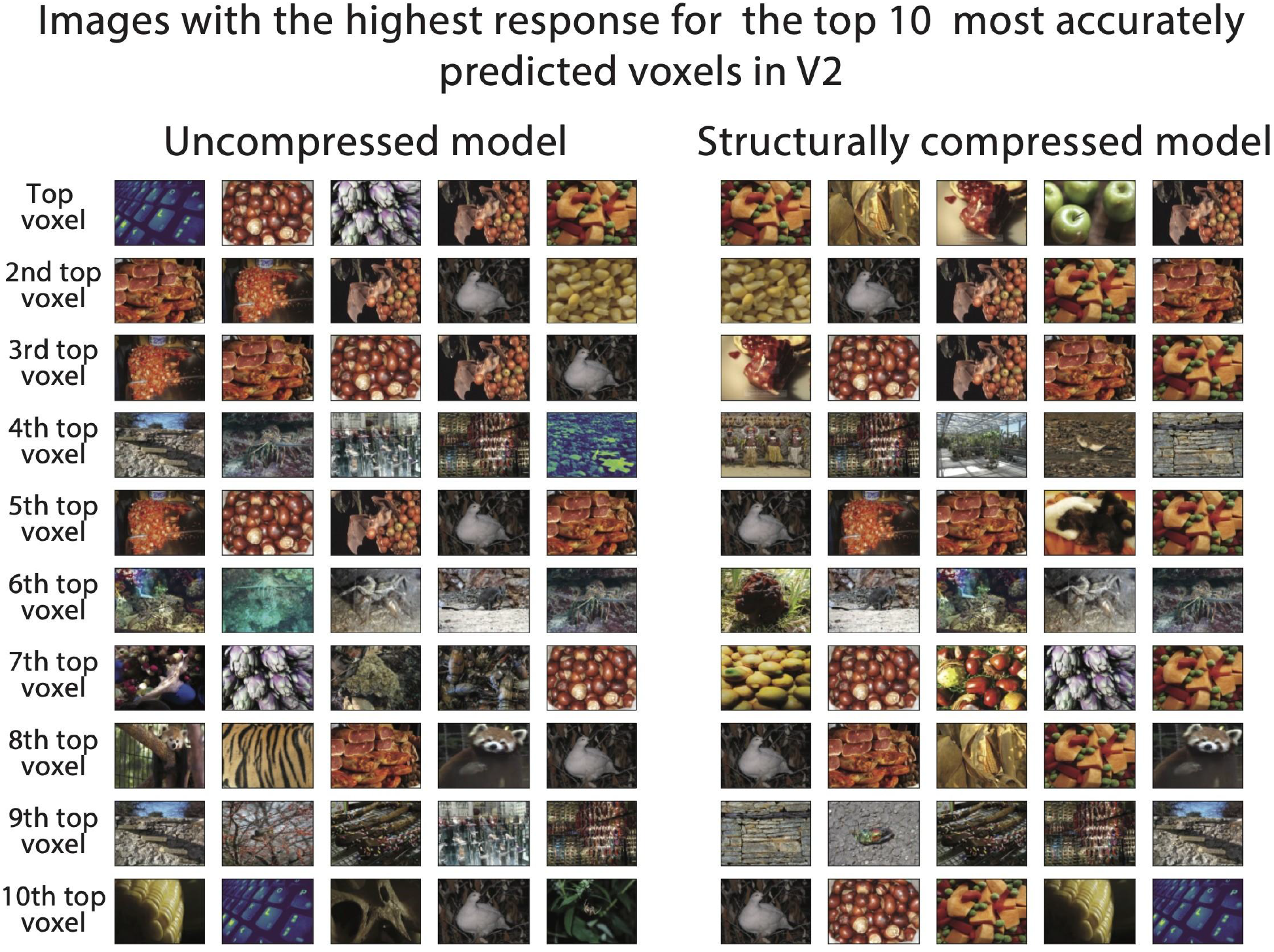
Comparing the performance of structurally-compressed and uncompressed models in V2. A. The five columns at left show the five images that the uncompressed model predicts will most increase activity for voxels in the V2 area, and the five columns at right show the five images that the structurally-compressed model predicts will most increase activity for voxels in the V2 area.

**Figure A10:**
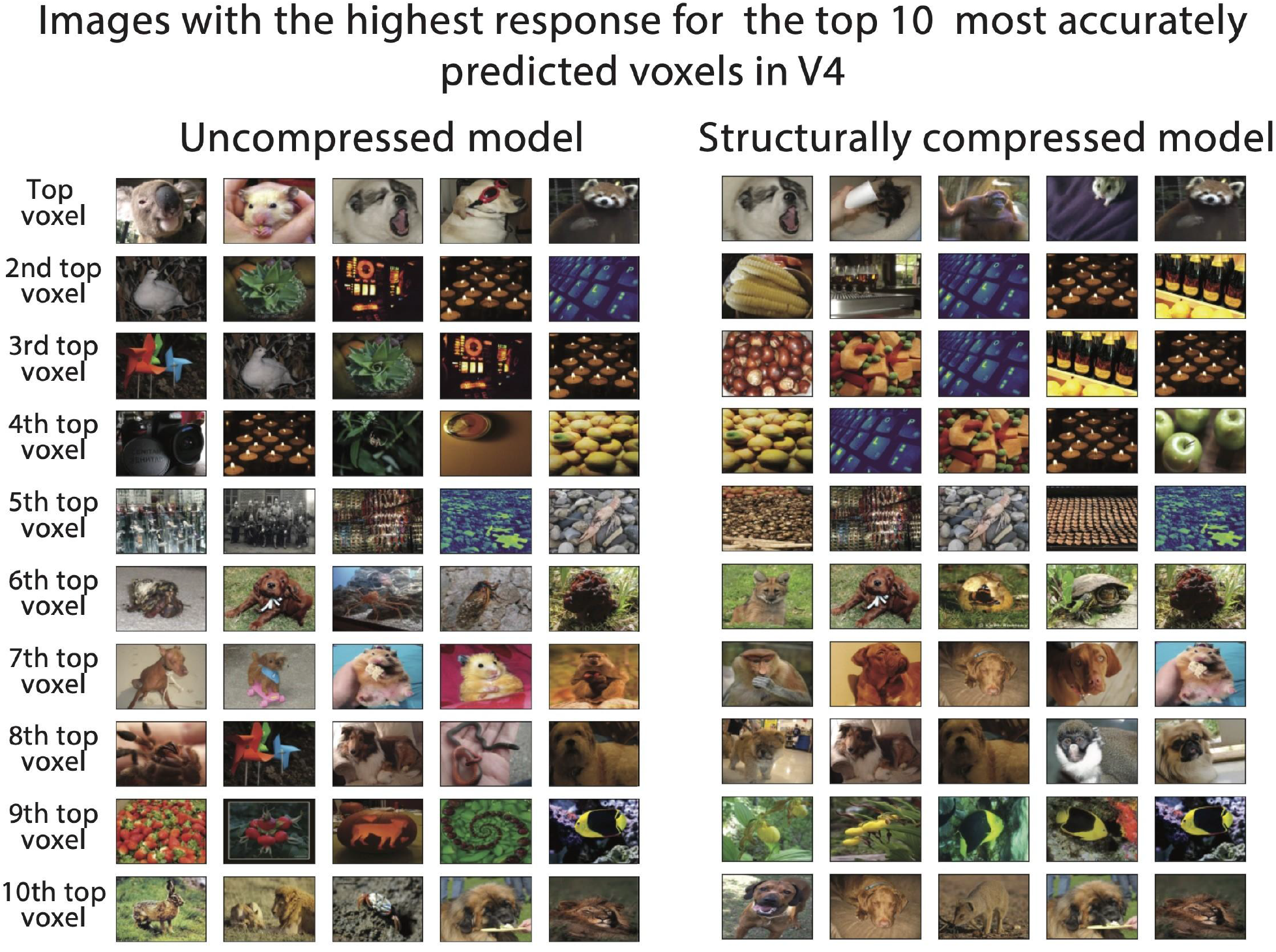
Comparing the performance of structurally-compressed and uncompressed models in V4. A. The five columns at left show the five images that the uncompressed model predicts will most increase activity for voxels in the V4 area, and the five columns at right show the five images that the structurally-compressed model predicts will most increase activity for voxels in the V4 area.

**Figure A11:**
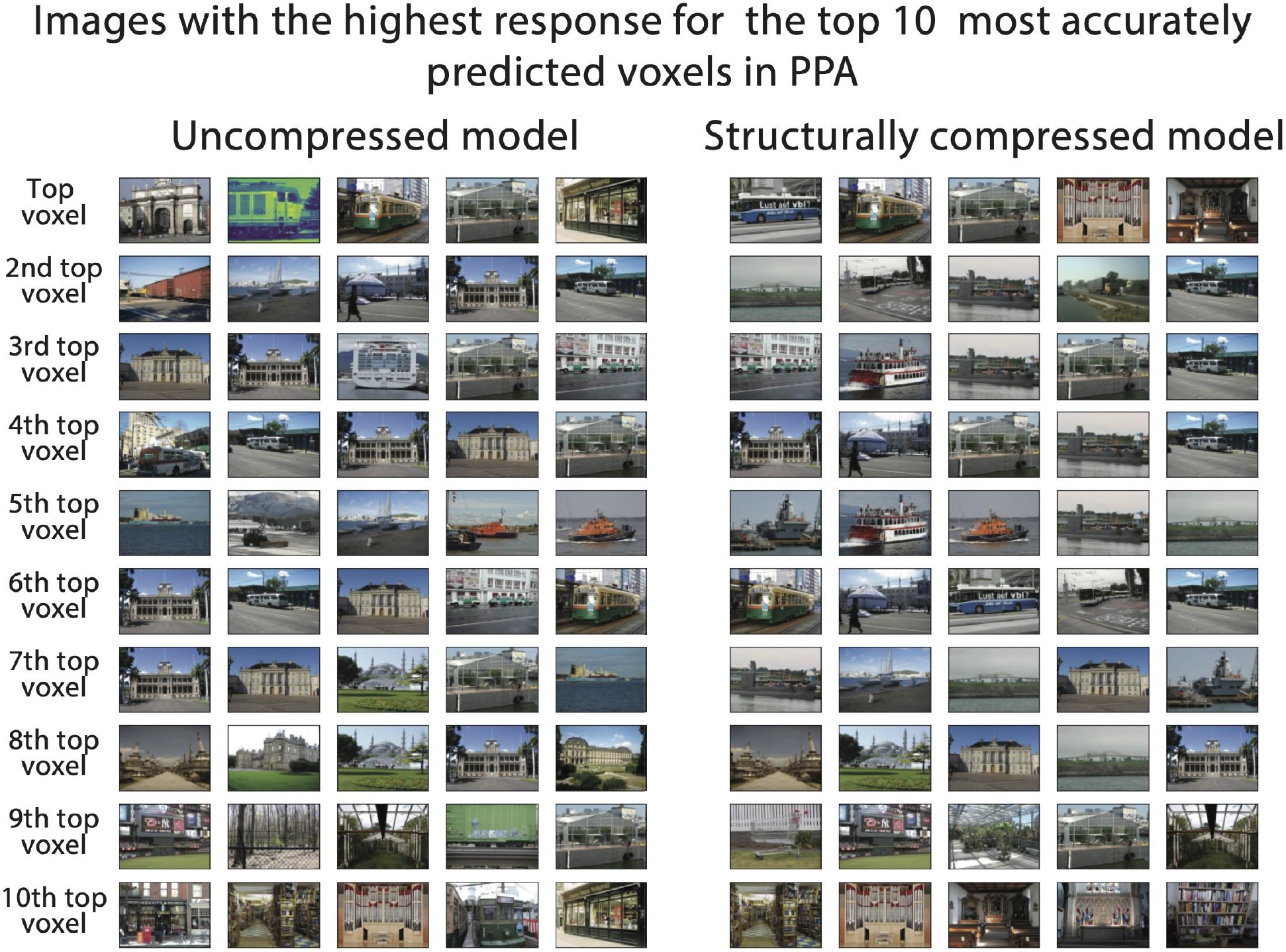
Comparing the performance of structurally-compressed and uncompressed models in PPA. A. The five columns at left show the five images that the uncompressed model predicts will most increase activity for voxels in the PPA area, and the five columns at right show the five images that the structurally-compressed model predicts will most increase activity for voxels in the PPA area.

**Figure A12:**
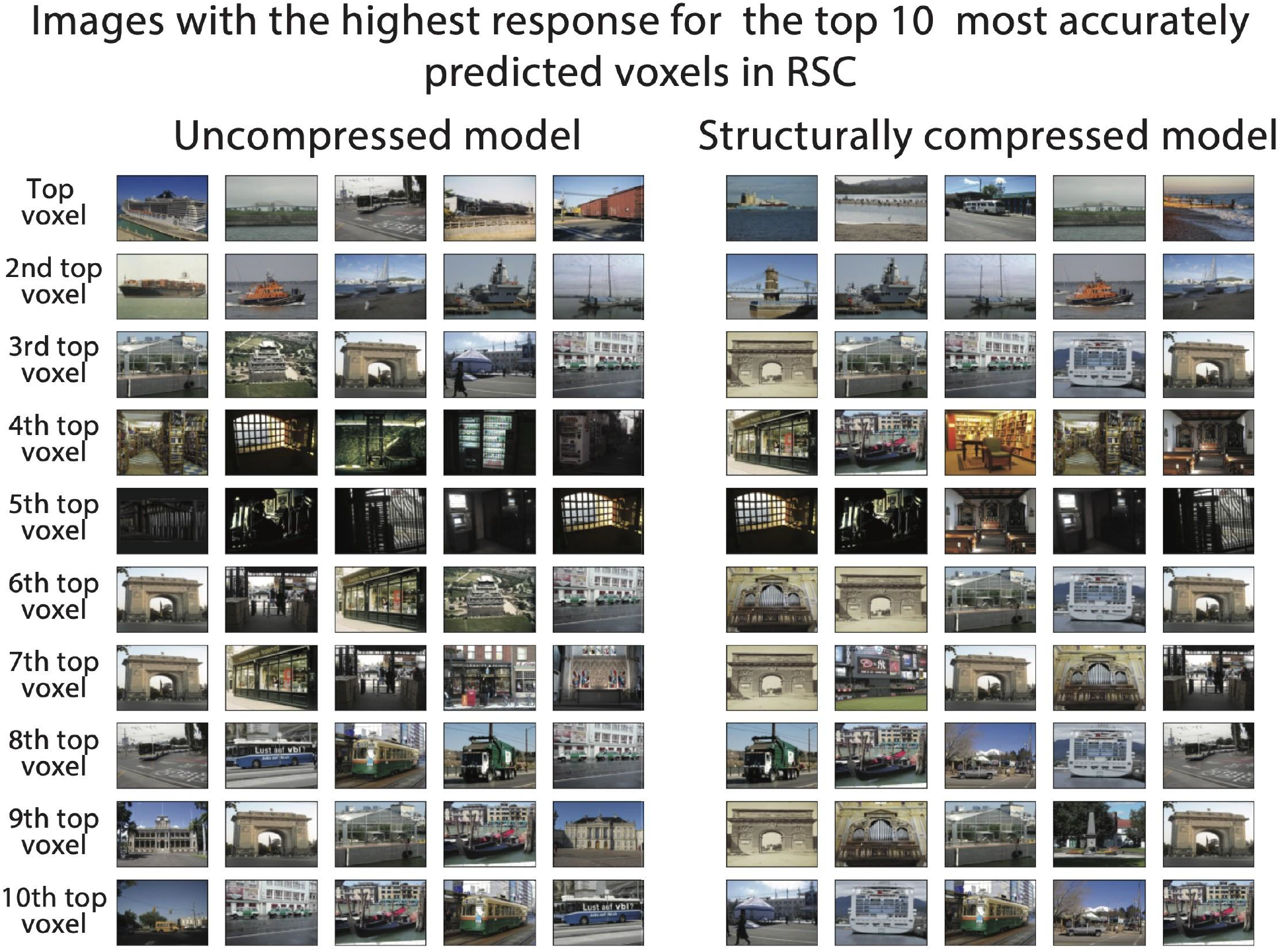
Comparing the performance of structurally-compressed and uncompressed models in RSC. A. The five columns at left show the five images that the uncompressed model predicts will most increase activity for voxels in the RSC area, and the five columns at right show the five images that the structurally-compressed model predicts will most increase activity for voxels in the RSC area.

